# *Pten* and *Dicer1* loss causes poorly-differentiated endometrial adenocarcinoma in mice

**DOI:** 10.1101/2020.03.12.989087

**Authors:** Xiyin Wang, Jillian R. H. Wendel, Robert E. Emerson, Russell R. Broaddus, Chad Creighton, Douglas B. Rusch, Aaron Buechlein, Francesco J. DeMayo, John P. Lydon, Shannon M. Hawkins

## Abstract

Endometrial cancer remains the most common gynecological malignancy in the United States. While the loss of the tumor suppressor, PTEN (phosphatase and tensin homolog), is well studied in endometrial cancer, recent studies suggest that *DICER1*, the endoribonuclease responsible for miRNA genesis, also plays a significant role in endometrial adenocarcinoma. In an endometrial adenocarcinoma mouse model, which has a conditional uterine deletion of *Pten, Dicer1* was also conditionally deleted. Conditional uterine deletion of *Dicer1* and *Pten* resulted in high-penetrance, poorly-differentiated endometrial adenocarcinomas. Poorly-differentiated endometrial adenocarcinomas expressed known markers of clear-cell adenocarcinoma, including Napsin A and HNF1B (hepatocyte nuclear factor 1 homeobox B). Adenocarcinomas were hormone-independent, and treatment with long-term progesterone did not mitigate poorly-differentiated adenocarcinoma, nor did it affect adnexal metastasis. Transcriptomic analyses of uteri or Ishikawa cells with deletion of *DICER1* revealed unique transcriptomic profiles and global downregulation of miRNAs. Integration of downregulated miRNAs with upregulated mRNA targets revealed deregulated let-7 and miR-16 target genes, similar to published human *DICER1*-mutant endometrial cancers from TCGA (The Cancer Genome Atlas). Importantly, these miRNA-target genes, involved in ephrin-receptor and transforming growth factor-beta signaling, represent potential clinical targets for rare, yet deadly, poorly-differentiated endometrial adenocarcinomas in women. This mouse model represents poorly-differentiated endometrial adenocarcinoma and will allow for the discovery of novel mechanisms of hormone-independent endometrial adenocarcinoma from atrophic endometrium.

**Significance Statement:** Endometrial cancer is one of the few cancers with an increasing death rate in the United States. The most significant risk factor associated with death is high tumor grade, which occurs most frequently in postmenopausal women, where it develops within an atrophic endometrium. Here, we present a mouse model with conditional deletion of *Dicer1*, a key enzyme in miRNA genesis, and *Pten*, a tumor suppressor, that develops poorly-differentiated, steroid hormone-independent, endometrial adenocarcinoma with adnexal metastasis. These high-grade adenocarcinomas develop from an atrophic endometrium and share molecular features with *DICER1*-mutant human endometrial adenocarcinomas. We anticipate that this preclinical model represents a move toward the discovery of novel mechanisms of hormone-independent development of endometrial adenocarcinoma from atrophic endometrium.

## Main Text

### Introduction

Endometrial cancer is the most common gynecologic malignancy in the United States and will account for 65,620 new cases and 12,590 deaths in 2020 (1). Well-differentiated [Fédération Internationale de Gynécologie Obstétrique (FIGO), grade 1] endometrioid endometrial adenocarcinoma represents the most common histology and has a 5-year survival for all stages that approaches 83% (2). High-risk histologic endometrial adenocarcinoma, such as poorly-differentiated (FIGO grade 3) endometrioid, serous, and clear-cell adenocarcinoma, encompasses 15% of endometrial cancer cases but accounts for nearly 50% of deaths (3, 4). High tumor grade is the most significant risk factor for disease recurrence and subsequent death (5, 6).

In endometrial cancer, DICER1 is a prominent cancer-driver gene (7). Hotspot biallelic mutations in *DICER1*, the endoribonuclease responsible for miRNA genesis, are enriched in endometrial adenocarcinomas over other cancers profiled in both TCGA (The Cancer Genome Atlas) PanCancer and MSK-IMPACT (Memorial Sloan-Kettering Integrated Mutation Profiling and Actionable Cancer Targets) studies (8). DICER1 contains two RNase III domains – RNase IIIa and RNase IIIb. These domains function to dice the precursor stem-loop miRNA molecule into two separate single-stranded mature miRNA molecules (9, 10). *In vitro* studies showed that hotspot mutations in the RNase IIIb domain resulted in altered miRNA processing (11–15). Independent studies not only validated the effects of hotspot biallelic mutations in the RNase IIIb domain resulting in miRNA biogenesis defects in human endometrial cancers but also showed that hotspot mutations in the RNase IIIa domain had similar functional effects in human endometrial cancers (8). Further, downregulation of *DICER1* expression was associated with features of more aggressive endometrial adenocarcinoma, including myometrial invasion, high FIGO grade tumors, and disease recurrence (16–18).

Uterine-specific deletion of *Dicer1* using *Pgr^Cre/+;^ Dicer1^flox/flox^* mice exhibited a uterine phenotype that is consistent with atrophic endometrium from postmenopausal women, containing a benign single layer of luminal epithelium, lack of glandular epithelium, and thin to zero endometrial stroma (19). Loss or mutation of the tumor suppressor, phosphatase and tensin homolog (*PTEN*), occurs in more than 80% of endometrial adenocarcinomas (20). Mice with conditional deletion of *Pten* in the uterus (*Pgr^Cre/+^; Pten^flox/flox^*) have high-penetrance (88.9%) well-differentiated endometrial adenocarcinoma by 90 days (21). Because global downregulation of *DICER1* expression was clinically associated with high FIGO grade endometrial cancer in women (16), we hypothesized that deletion of *Dicer1* with deletion of *Pten* would result in high FIGO grade endometrial adenocarcinomas.

In this study, we present data that underscores the impact of appropriate *Dicer1* function in endometrial adenocarcinoma. Importantly, we describe the development of hormone-independent, poorly-differentiated adenocarcinoma with adnexal metastasis that arises from an atrophic endometrium. High-fidelity mouse models of poorly-differentiated endometrial adenocarcinomas have not been described. Further, xenograft models have shown limited success for high-risk histologic endometrial adenocarcinoma types (22). Our findings indicate a relationship between appropriate miRNA expression, cell cycle control, and cell signaling pathways in the development of poorly-differentiated endometrial adenocarcinoma. Elucidating the role of *DICER1* and miRNA dysregulation in poorly-differentiated endometrial adenocarcinoma will uncover novel mechanisms that can be targeted for therapies in women.

## Results

### Bulky endometrial tumors in adult mice with loss of two alleles of *Dicer1*

To study the role of *Dicer1* in endometrial cancer, *Dicer1* was conditionally deleted in *Pgr^Cre/+;^ Pten^flox/flox^* mice. *Pgr^+/+^* (control), *Pten* cKO [*Pten* conditional knockout (*Pgr^cre/+^*; *Pten^flox/flox^; Dicer1^+/+^*)], and dcKO [double conditional knockout (*Pgr^cre/+^*; *Pten^flox/flox^; Dicer1^flox/flox^*)] were generated. *Pten-Dicer* het [*Pten*-*Dicer* heterozygous (*Pgr^cre/+^*; *Pten^flox/flox^; Dicer1^flox/+^*)] were studied in parallel, but they were similar to *Pten* cKO (Supplementary Figure S1 and Supplementary Table S1). Examination of the reproductive tract of 6-month-old female mice revealed bulky endometrial tumors in both *Pten* cKO and dcKO mice (Figure 1A). Uteri from *Pgr^+/+^* mice were normal with grossly normal adnexa. Uteri from *Pten* cKO mice revealed large tortuous uteri with bulky nodules of solid tumors protruding onto the surface, but homogeneous in gross morphology along both uterine horns. Uteri from dcKO mice were significantly smaller but contained areas of solid, bulky tumors. Histological analyses revealed adenocarcinoma invading through the myometrium of the uterus and into the adnexa, but without gross metastatic disease outside the female reproductive tract (Supplementary Figure S2-3). Due to morbidity, only a small number of mice at this time point were examined (Supplementary Tables S2).

**Figure 1.**
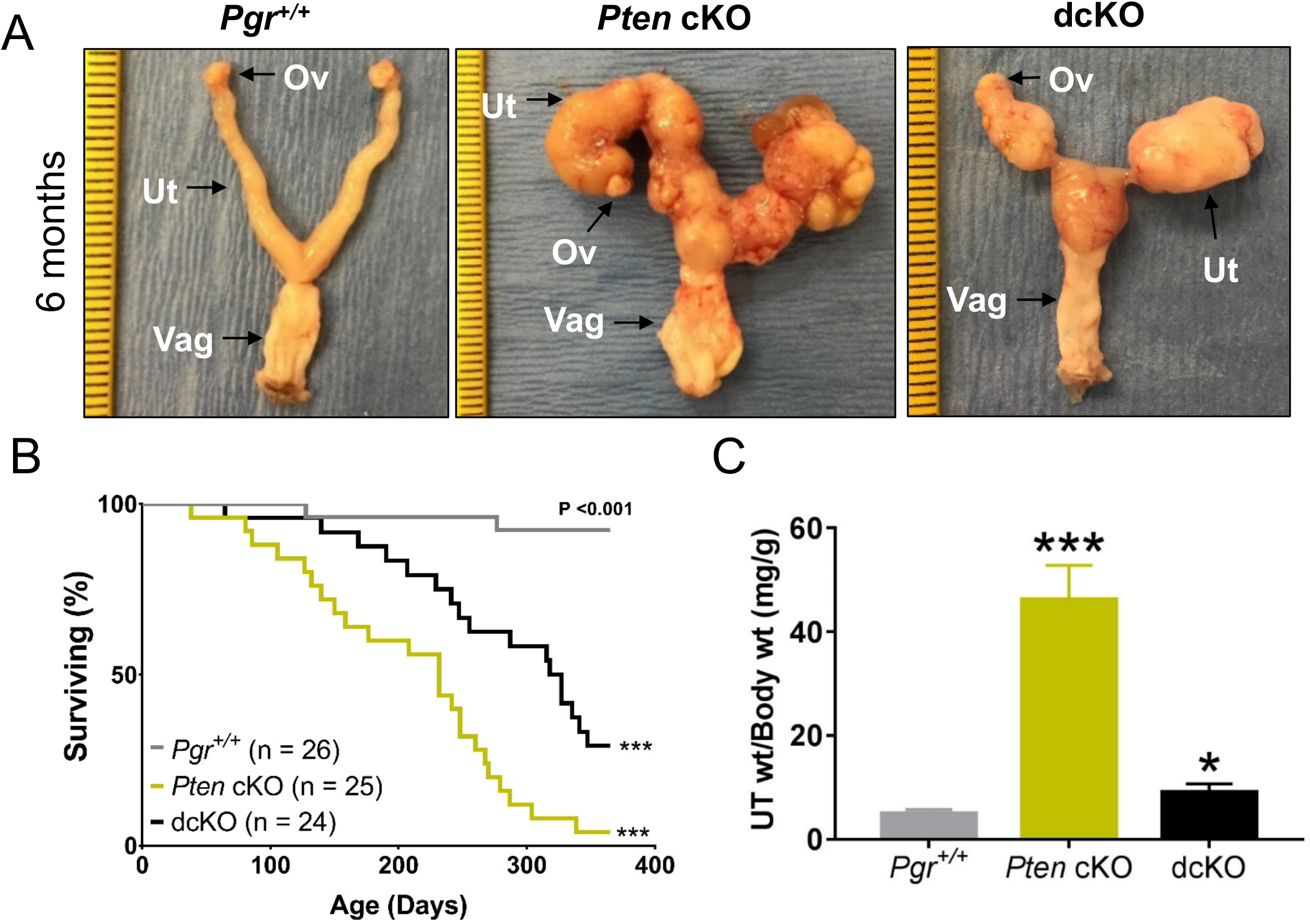
Unique morphology and poor survival with deletion of *Dicer1*. (A) Postnatal mice at 6 months were dissected, and female reproductive tracts were examined grossly. Tumor burden was seen in both *Pten* cKO and dcKO mice. Ov, ovary; Ut, uterus; Vag, vagina. (Ruler, tick mark indicates 1mm, but dcKO photo is further zoomed.) (B) Kaplan-Meier survival curves were analyzed by log-rank (Mantel-Cox) pairwise comparison with Bonferroni correction. Both *Pten* cKO and dcKO mice have decreased survival compared to control mice. ***, *P* < 0.001. (C) At 12 weeks, uteri were weighed and normalized to body weight. Both *Pten* cKO and dcKO mice had larger uteri than *Pgr^+/+^*. Mean ± SEM; *, *P* < 0.05; ***, *P* < 0.001; multiple *t*-test; *n* > 8 per group.

*Pten* cKO and dcKO mice showed decreased survival compared to *Pgr^+/+^* (Figure 1B). *Pten* cKO had a median survival of 240 days, similar to published results (21). The median survival for dcKO mice was 327 days. Many mice had necrotic tumors throughout the reproductive tract, making an accurate assessment of tumor histology impossible (Supplementary Tables S2).

### Poorly-differentiated adenocarcinoma in dcKO mice

By 12 weeks, dcKO uteri were nearly twice as large as *Pgr^+/+^* but remained significantly smaller than *Pten* cKO (Figure 1C). Uterine morphology, survival studies, and uterine weight suggested distinct phenotypes between *Pten* cKO and dcKO uteri. At 12 weeks, *Pgr^+/+^* uteri showed normal histology (Figure 2A). Similar to published (21), a majority of the *Pten* cKO uteri contained well-differentiated adenocarcinoma, with glands in columnar architecture and nuclear pseudostratification but with retained polarity in many of the nuclei, mild nuclear atypia, and little or no solid pattern. Endometrial adenocarcinoma was defined as confluent growth of the endometrium, with cribriform pattern and/or expansion of the endometrium into the endometrial stroma, or invasion into the surrounding myometrium (23). FIGO grading was used (24). With deletion of *Dicer1*, the frequency of poorly-differentiated adenocarcinoma increased significantly (χ2 = 12.8; *P* = 0.00035). Most of the dcKO mice exhibited poorly-differentiated adenocarcinoma (Supplementary Table S2), as evidenced by poorly formed malignant glands, solid microscopic architecture, or tubulocystic architecture (Figure 2A). Epithelial cells were stained with cytokeratin-8 and myometrium with smooth muscle actin (Supplementary Figure S4). Myometrial invasion was defined as tumor between the two layers of the myometrium. Similar to published results (21), 85% of tumors from *Pten* cKO uteri exhibited invasion through the myometrium. All dcKO adenocarcinomas at 12 weeks showed myometrial invasion (Supplementary Table S2).

**Figure 2.**
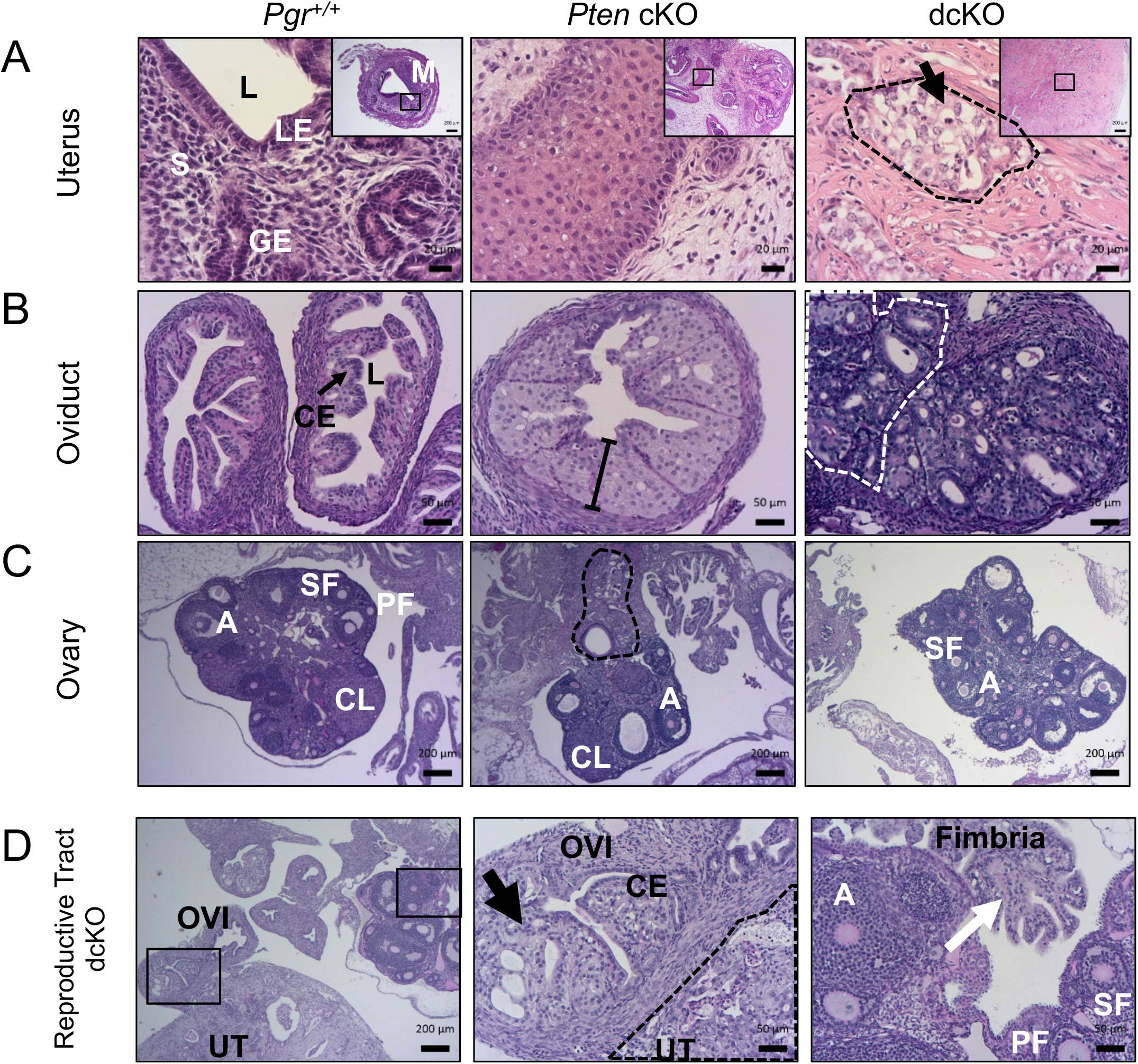
Invasive adenocarcinoma with metastasis to the adnexa. (A) *Pgr^+/+^* uteri showed normal luminal epithelium (LE) surrounding a lumen (L), with glandular epithelium (GE) and endometrial stroma (S), and two layers of smooth muscle in the myometrium (M). *Pten* cKO uteri exhibited well-differentiated adenocarcinoma. Uteri from dcKO mice exhibited poorly-differentiated adenocarcinoma (black dashed lines) with pale cytoplasm (black arrow). H&E. (Scale bars, low-power, 200μm; high-power, 20 μm.) (B) *Pgr^+/+^* oviducts contained a single layer of columnar epithelium (CE, arrow) surrounding a lumen (L), surrounded by a layer of smooth muscle. *Pten* cKO oviduct showed atypical epithelium as evidenced by >1 cell thickness epithelium (line). The dcKO oviduct showed adenocarcinoma within the oviduct as well as next to the oviductal lumen (grey dashed lines). (Scale bars, 50 μm.) (C) *Pgr^+/+^* ovary showed normal histology with follicles in each stage of follicular development. There were primary (PF), secondary (SF), and antral (A) follicles and corpus luteum (CL). *Pten* cKO ovaries had normal appearing follicles and well-differentiated adenocarcinoma (black dashed lines) that appeared to be invading from outside the ovary. Ovaries from dcKO mice did not frequently show adenocarcinoma close to the ovary. (Scale bars, 200 μm.) (D) Adult dcKO female reproductive tract showed direct invasion from moderately-differentiated adenocarcinoma in the uterus (UT) to the oviduct (OVI). Inset shows normal single layer of columnar epithelium (CE) in the oviduct next to adenocarcinoma (black arrow). Distal fimbria (white arrow) was normal. Ovary showed normal follicles, including primary (PF), secondary (SF), and antral (A) follicles. (Scale bars, low-power, 200 μm; high-power, 50 μm.)

### Malignant metastatic adnexal pathology

*Pgr^Cre/+^* mice have homologous recombination of floxed alleles within the columnar epithelial cells of the oviduct and the corpus luteum of the ovary when stimulated with gonadotropins (25). Older mice of all genotypes (*i.e.*, survival and 6-month old) exhibited adenocarcinoma of the uterus, oviduct, and ovaries. However, the origin of the tumor, driven by *Cre*-recombination events in the oviduct and/or ovary versus metastatic disease from the uterus, was unable to be discerned. Further, high-grade serous ovarian cancer develops in the oviduct of mice with mesenchymal deletion of both *Pten* and *Dicer1* (26). Moreover, activation of WNT/β-catenin through deletion of adenomatous polyposis coli (*Apc*) with *Pgr^Cre/+^* showed frequent endometrioid ovarian cancer that began in the epithelial cells of the oviduct (27). The oviducts and ovaries were carefully examined for any signs of cancer-associated histological changes. *Pten* cKO oviducts frequently showed epithelial atypia in the oviducts defined as nuclear stratification (>1 cell thickness epithelium, loss of nuclear polarity with nuclei not lined up near the interface with stroma) and nuclear atypia in the form of enlarged, rounded nuclei (Figure 2B). Adenocarcinoma was infrequently discovered in the oviducts of *Pten* cKO mice. When adenocarcinoma was discovered, the histology was similar to uterine adenocarcinoma and present inside and outside the oviduct, consistent with metastatic disease (Supplementary Table S2). Adenocarcinoma was found in nearly 50% of the oviducts of dcKO mice. The histology was similar to uterine histology, adenocarcinoma was located on the inside and outside of normal oviducts (Figure 2B), and there were no mice with adnexal tumors without uterine tumors, consistent with metastatic spread. *Pten* cKO ovaries showed evidence of both normal follicular development and metastatic disease, as cancer appeared to have invaded at the ovarian hilum (Figure 2C). Ovaries of dcKO mice were most frequently normal (Figure 2C-D). Examination of the reproductive tracts of dcKO mice showed evidence of direct extension from the uterus to the oviduct with normal, cancer-free, distal fimbria and ovary (Figure 2D). Evidence of metastatic disease in the adnexa (*i.e.*, oviduct or ovary) was observed in approximately 50% of *Pten* cKO and dcKO mice (Supplementary Table S2).

### Poorly-differentiated dcKO tumors expressed clear-cell adenocarcinoma markers

Epithelial cells from dcKO uterine tumors exhibited large, often rounded, and hyperchromatic nuclei and pale-staining to clear cytoplasm (Figure 2A and Supplementary Figure S2). To better describe the poorly-differentiated adenocarcinomas with clear cytoplasm in dcKO mice, immunohistochemical markers of clear-cell adenocarcinoma, Napsin A and hepatocyte nuclear factor 1 homeobox B (HNF1B) (28, 29), were evaluated. Poorly-differentiated adenocarcinomas from dcKO mice exhibited malignant epithelial cells with high-frequency, high-intensity cytoplasmic Napsin A staining. Malignant epithelial cells from dcKO mice more frequently stained positive for nuclear HNF1B compared to *Pten* cKO tumors (Figure 3 and Supplementary Figure S5).

**Figure 3.**
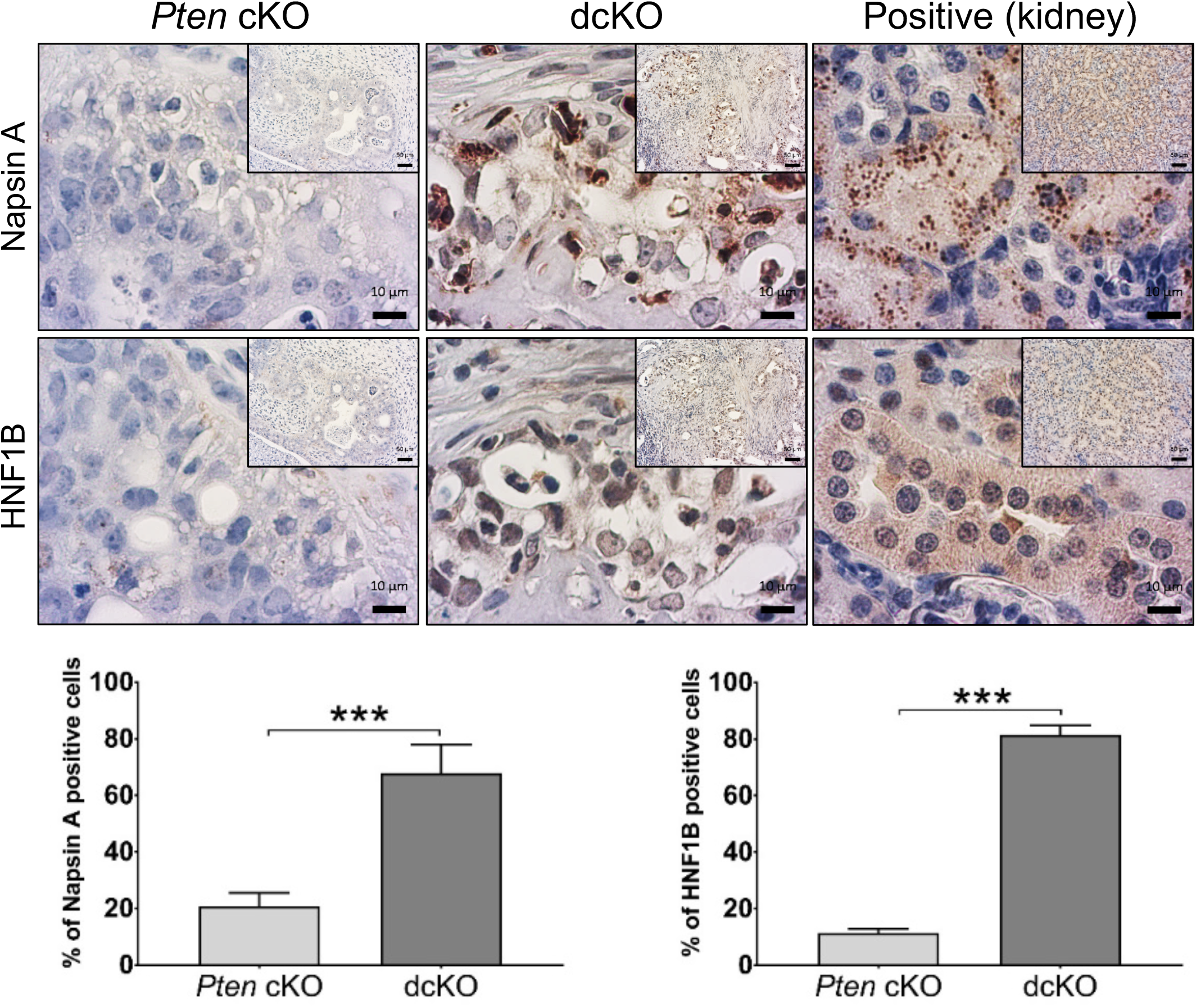
Poorly-differentiated adenocarcinomas from 12-week-old dcKO mice expressed clear-cell adenocarcinoma markers. Napsin A staining was cytoplasmic, and HNF1B staining was nuclear. Quantification of the frequency of positive epithelial cells. Kidney is shown as positive control. Mean expression scores were compared using 2-tailed Student’s *t*-test. Average scoring ± SEM is shown; ***; *P*<0.001; *n* = 6. (Scale bars, low-power, 50 μm; high-power, 10 μm.)

### No effect on the frequency of poorly-differentiated adenocarcinoma histology with steroid hormone depletion and/or progesterone treatment

Recent work (30) showed that genetic changes in *Pten*, phosphatidylinositol-4,5-bisphosphate 3-kinase catalytic subunit alpha (*Pik3ca*), and catenin beta 1 (*Ctnnb1*) within the mouse uterine epithelium were not sufficient for malignant transformation. However, ovarian insufficiency was required for malignant transformation of those “triple” mutant endometrial epithelial cells (30). To determine the effects of ovarian insufficiency in dcKO mice, both ovaries were removed, and uterine weight and histology were assessed. Gross uterine weight was lower in ovariectomized (ovex) mice compared to intact mice (Supplementary Figure S6A). However, the frequency of poorly-differentiated adenocarcinoma was not decreased in dcKO mice (Supplementary Table S2).

Studies showed that triple mutant (*Pten, Pik3ca, Ctnnb1*) endometrial epithelial cells transformed with steroid hormone depletion in endometrial adenocarcinoma were responsive to progesterone (30). To examine the effects of long-term progesterone therapy, mice underwent removal of ovaries at six weeks and were randomized to either placebo or progesterone pellets at eight weeks. Treatment lasted 60 days. Treatment with progesterone did not alter the frequency of poorly-differentiated adenocarcinomas in ovex *Pten* cKO or dcKO mice (Supplementary Table S2). Carefully selected women with well-differentiated, early-stage endometrioid adenocarcinoma can be treated with progesterone to preserve fertility options (22). Intact mice were randomized at six weeks to either placebo or progesterone pellet for 60 days. Treatment with progesterone did not significantly alter the frequency of poorly-differentiated endometrial adenocarcinomas in intact mice (Supplementary Table S2). Treatment of intact mice with long-term progesterone decreased the rate of metastatic adenocarcinoma to the adnexa, but it was only statistically significantly decreased for *Pten* cKO mice (Supplementary Figure S6B; Supplementary Table S2).

### Unique transcriptomic profile with loss of two alleles of *Dicer1*

To create an *in vitro* human model, CRISPR-Cas9 was used to delete *DICER1* in Ishikawa cells, as they are *PTEN* mutant (22). Western blot confirmed knockdown or knockout of DICER1 (Figure 4A). *DICER1* deletion in Ishikawa cells led to decreased cellular proliferation and reduced colony formation in soft agar (Figure 4B-C), consistent with the smaller-sized uterine tumors in dcKO mice (Figure 1C). Poly-A RNA sequencing (RNA-seq) was performed. Principal Component (PC) analysis of the RNA-seq data showed distinct clustering of each sample. *DICER1^+/+^, DICER1^+/−^*, and *DICER1^−/−^* differed significantly in PC2, while *DICER1^+/+^* and *DICER1^+/−^* were most similar in PC1 (Figure 4D). Determination of differentially expressed genes between *DICER1^+/+^* and *DICER1^−/−^* revealed 2444 unique protein-coding genes with log2fold change >|1| significantly dysregulated [false discovery rate (FDR)<0.05, Supplementary Table S3], 1154 genes upregulated, and 1290 genes downregulated (Supplementary Figure S7A).

**Figure 4.**
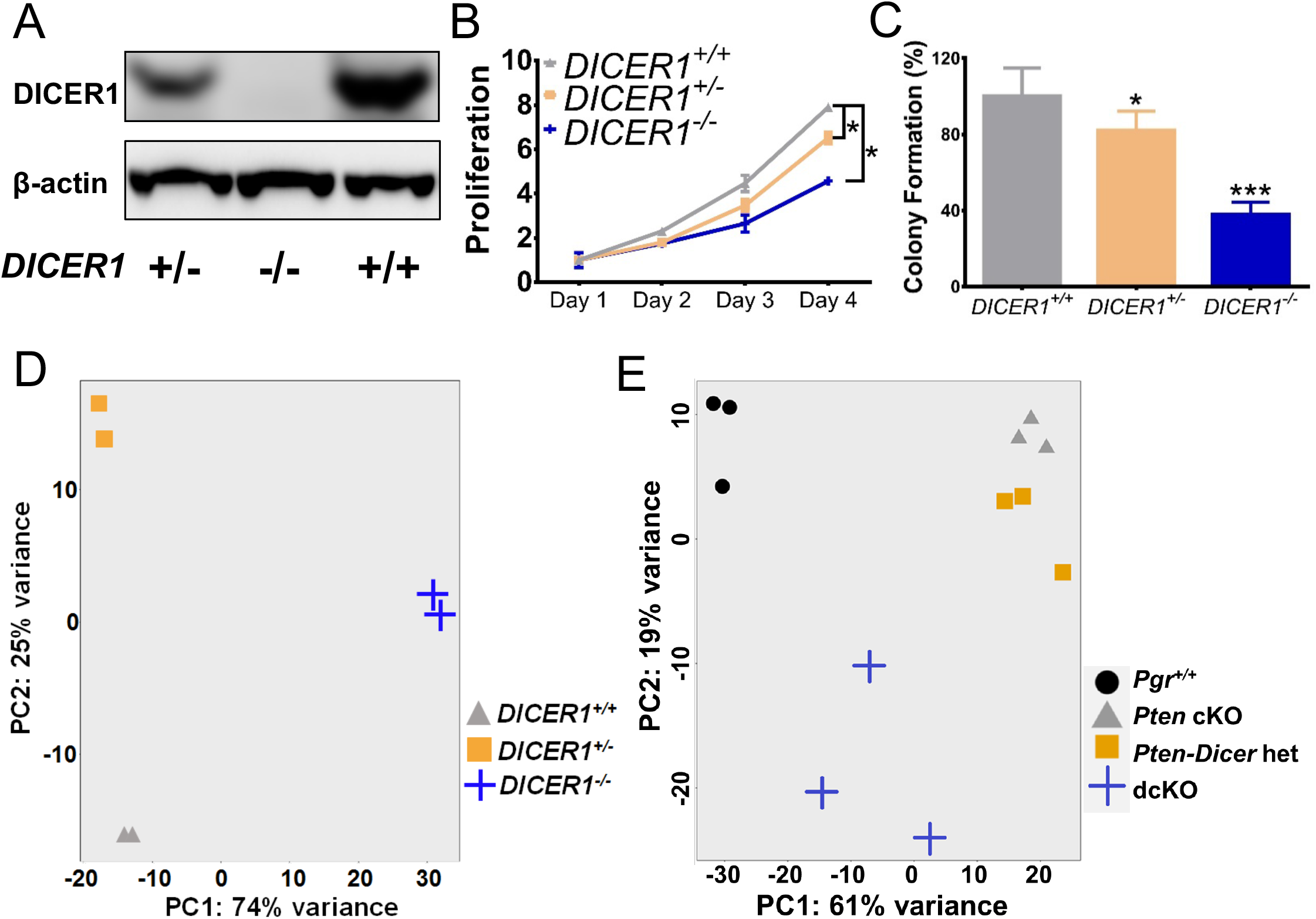
Unique cellular characteristics and transcriptomic profile with loss of two alleles of *DICER1*. (A) Western blot showed decreased DICER1 protein levels in *DICER1^+/−^* and complete loss of DICER1 protein in *DICER1^−/−^* cells. Both *DICER1^+/−^* and *DICER1^−/−^* cells showed a significant decrease in (B) proliferation and (C) colony formation relative to *DICER1^+/+^*. *, *P*<0.05, ***, *P*<0.001. (D) Principal component analysis revealed distinct clustering of human samples in *DICER1^+/+^* (triangles), *DICER1^+/−^* (squares), and *DICER1^−/−^* (plus sign). (E) Principal component analysis for mouse showed close clustering of *Pten* cKO (triangles) and *Pten-Dicer* het (squares) but distinct clustering of *Pgr^+/+^* (circles) and dcKO (plus sign).

Poly-A RNA sequencing on mouse uteri was then performed. Three-week uteri were selected because: 1) *Pgr^Cre/+^-*mediated deletion occurs primarily in the luminal and glandular epithelium (25); 2) the limited molecular contributions of steroid hormones in pre-adolescent female mice; 3) the ability to explore early molecular contributions; and 4) the similarity of cell populations (*i.e.*, epithelium, stroma, and myometrium) across genotypes. As early as three weeks, *Pten* cKO and dcKO uteri showed adenocarcinoma (Supplementary Table S2) with activation of phospho-AKT limited to the epithelium (Supplementary Figure S8). PC analysis showed distinct clustering of each genotype. However, *Pten* cKO and *Pten-Dicer* het mRNA profiles were most similar to each other and only varied slightly in PC2. Both dcKO and *Pgr^+/+^* varied significantly in PC1, and dcKO varies in PC2 (Figure 4E). Because all experimental mice have a deletion of *Pten*, transcriptomic profiles were compared to *Pten* cKO instead of *Pgr^+/+^*. Determination of differentially expressed genes between *Pten-Dicer* het and *Pten* cKO revealed 13 unique transcripts (FDR<0.05, Supplemental Table S3). Determination of differentially expressed genes between dcKO and *Pten* cKO showed 1635 unique protein-coding genes with log2fold change >ǀ1ǀ significantly dysregulated (FDR<0.01, Supplementary Figure S7B), 1059 genes upregulated, and 576 genes downregulated (Supplementary Table S3).

Gene set enrichment analysis with estrogen-responsive gene sets (31) showed no significant overlap with mouse dcKO or human Ishikawa *DICER1^−/−^* transcriptomic profile (Supplementary Table S4). The genes downregulated in mouse dcKO showed a trend (*P*=0.087) towards enrichment in progesterone-responsive gene sets (32), but no significant enrichment in upregulated progesterone-responsive genes. Both up- and downregulated genes in the human Ishikawa *DICER1^−/−^* transcriptomic profile showed significant enrichment in progesterone responsive genes (Supplementary Table S4).

Using Web Gestalt (WEB-based Gene SeT AnaLysis Toolkit) (33) revealed that the upregulated human Ishikawa *DICER1^−/−^* transcriptomic profile was enriched in ephrin receptor signaling pathway genes (Supplementary Table S3). Studies have shown that high expression of EPH receptor A2 (*EPHA2)* in endometrial cancer was associated with high tumor grade, reduced survival, high microvessel density, and high Ki67 expression (34, 35). QPCR of *DICER1^−/−^* cells showed that *EPHA2* was 7.6-fold upregulated. Other EPH receptors, including EPH receptor A7 (*EPHA7*) and EPH receptor B2 (*EPHB2*), were 8-10-fold upregulated (Figure 5A). Further, both the non-canonical WNT signaling molecule, *WNT5B*, and p21 (RAC1) activated kinase 3 (*PAK3*) downstream effectors of EPHA2 (36, 37), were 9.6- and 9.3-fold upregulated. LIM kinase 2 (*LIMK2*), an essential molecule in p21 signal transduction and involved in castration-resistant prostate cancer (38), was similarly upregulated (Figure 5A). QPCR on dcKO uteri showed that EPH receptor B1 (*Ephb1*) was 3.5-fold upregulated, and *Pak3* was 5.3-fold upregulated. Cofilin 2 (*Cfl2*), a potential downstream effector of LIMK2 and PAK3 (39), was 2.7-fold upregulated (Figure 5B). Pathway analysis on the upregulated dcKO uteri transcriptomic profile revealed hedgehog signaling and prostaglandin synthesis genes were upregulated (Supplementary Table S3). QPCR of dcKO uteri showed Indian hedgehog signaling molecule (*Ihh*) 20-fold upregulated and patched 1 (*Ptch1*) and GLI family zinc finger 1 and 2 (*Gli1* and *Gli2*) were 5-fold upregulated (Figure 5C). High CFL2 expression was associated with poor survival in gastric cancer (40). Both high CFL2 and EPHA2 have been associated with increased proliferation in cancers (34, 35, 40). Examination of dcKO uteri showed increased expression of Ki67 compared to *Pten* cKO uteri (Figure 5D).

**Figure 5.**
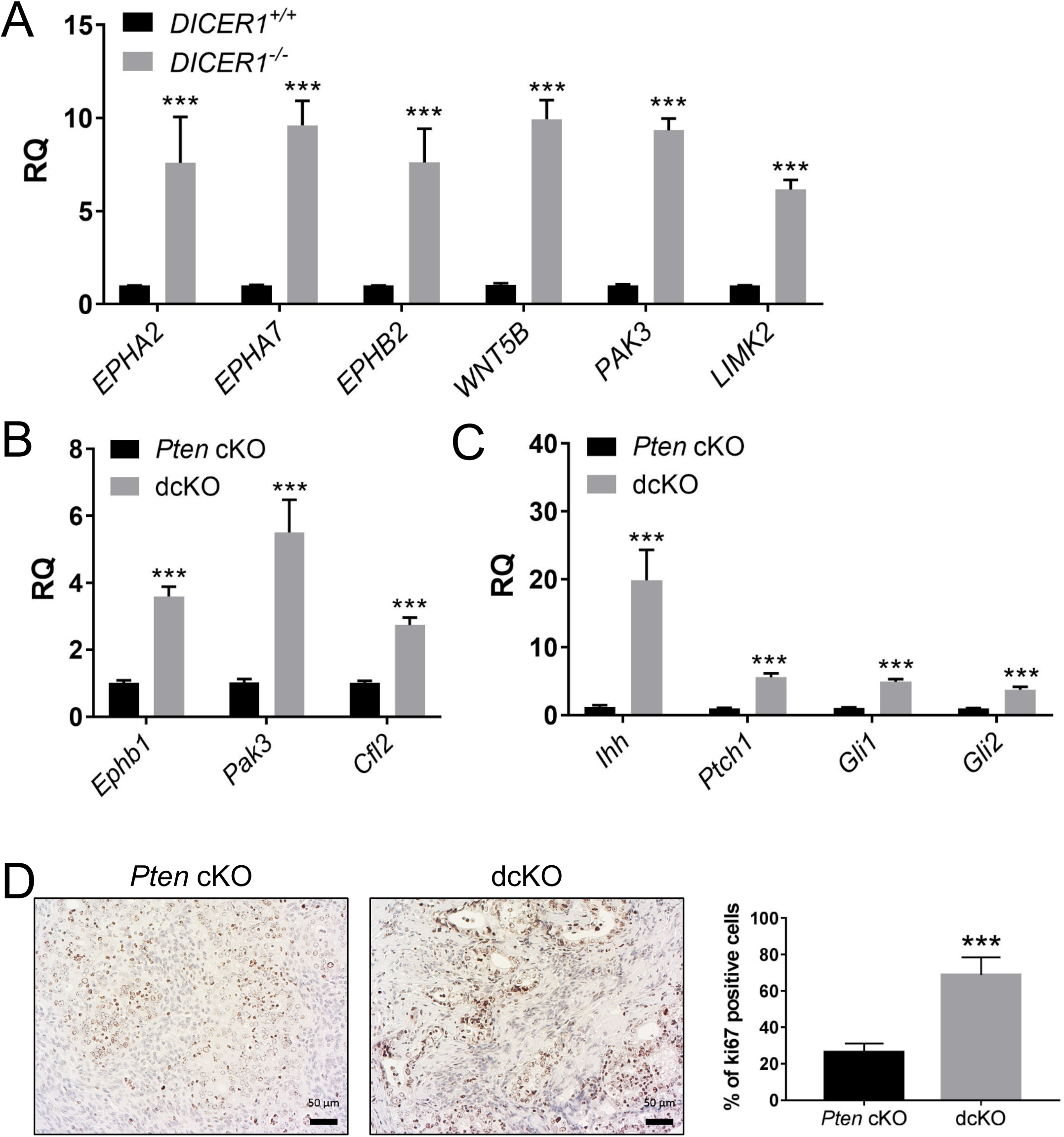
Dysregulation of ephrin receptor signaling genes with *DICER1* deletion. QPCR showed upregulation of ephrin receptor signaling genes in human *DICER1^−/−^* cells (A) and dcKO mice (B). QPCR showed upregulation of hedgehog signaling genes in dcKO uteri (C). Mean± SEM; two-tailed Student’s *t*-test; ***, *P* < 0.001; *n* = 6; RQ, relative quantity of each gene transcript relative to endogenous control gene *GAPHD* (human) and 18S (mouse) normalized to *DICER1^+/+^* (A) or *Pten* cKO (B-C). (D) Elevated Ki67 expression in dcKO compared *Pten* cKO. (Scale bars, 50 μm.) Mean± SEM; two-tailed Student’s *t*-test; ***, *P* < 0.001; *n* > 8.

To examine the effects of *DICER1* deletion on miRNA expression in endometrial cancer, small RNA sequencing was performed on the same samples as poly-A RNA sequencing, both mouse and human. Differentially expressed miRNAs (FDR<10%) in mouse dcKO and human Ishikawa *DICER1^−/−^* cells are shown in Supplementary Table S5. Human Ishikawa *DICER1^−/−^* cells showed 193, and dcKO uteri revealed 38 mature miRNAs downregulated. There were 22 mature miRNAs downregulated in common. MiRNAs target mRNAs for repression through binding to the 3’ untranslated region of target genes (41). Integration of upregulated miRNA-target genes with downregulated miRNAs from human Ishikawa *DICER1^−/−^* cells was performed using miRTarBase. MiRTarBase uses curated data from over 10,000 publications to make validated predictions on miRNA-target interactions (MTIs) (42, 43). Out of 1154 upregulated genes in human Ishikawa *DICER1^−/−^* cells, 885 were MTIs for the 193 downregulated miRNAs. Supplementary Table S5 lists the 7023 miRNA:mRNA MTIs mapped to 885 unique genes. Using Web Gestalt (33), the genes targeted by miRNAs were involved in the following pathways: microRNAs in cancer, glycosaminoglycan biosynthesis, ephrin-receptor signaling, and transforming growth factor-beta (TGFβ) signaling (Supplementary Table S5). Many TGFβ signaling genes were also dysregulated in dcKO uteri (Supplementary Table S5).

Human *DICER1*-mutant TCGA endometrial cancers were enriched in miRNA-target genes from five miRNA families: let-7-5p, miR-17-5p, miR-16-5p, miR-29-3p, and miR-101-3p (8). Mature miRNAs from each of these five miRNA families were significantly downregulated in the small RNA sequencing datasets from human Ishikawa *DICER1^−/−^* cells (Supplementary Table S5). Out of 1154 upregulated genes in human Ishikawa *DICER1^−/−^* cells, 347 unique genes were MTIs for these specific five *DICER1*-mutant family members. Pathways enriched included microRNAs in cancer, glycosaminoglycan biosynthesis, ephrin-receptor signaling, renal cell carcinoma, and TGFβ signaling (Supplementary Table S5).

Recent work has shown that dysregulation of TGFβ signaling in the mouse uterus results in endometrial cancer (44, 45). Members of the TGFβ signaling pathways were targeted by a number of dysregulated miRNAs. Let-7b-5p, miR-16-5p, and miR-26b-5p were mature miRNAs downregulated in both human Ishikawa *DICER1^−/−^* cells and dcKO uteri and predicted to target TGFβ signaling genes. By QPCR, *DICER1^−/−^* cells had significant downregulation of let-7b-5p, and let-7b-5p was 2-fold down-regulated in dcKO uteri. QPCR showed a 71-fold downregulation of miR-16-5p in human Ishikawa *DICER1^−/−^* cells, and 2.1-fold downregulation for miR-16-5p in mouse. QPCR showed 939-fold downregulation of miR-26b-5p in human and 1.7-fold downregulation in the dcKO uteri (Figure 6A-B). Other miRNAs validated by QPCR are shown in Supplementary Figure S9. These miRNAs were validated targets for the TGFβ signaling genes: cyclin-dependent kinase 6 (*CDK6*), bone morphogenetic protein receptor 1B (*BMPR1B*), and transforming growth factor-beta receptor 3 (*TGFBR3*, Supplementary Table S5). MiR-26b-5p was also a validated target for *CDK6*, transforming growth factor-beta 1 induced transcript 1 (*TGFB1I1*), and growth differentiation factor 10 (*GDF10*, Supplementary Table S5). QPCR showed higher than 6.5-fold upregulation of *CDK6* in human and 2.5-fold upregulation of *Cdk6* in the mouse. *BMPR1B* showed a greater than 14-fold upregulation in *DICER1^−/−^* cells. *TGFBR3* showed 6.5-fold upregulation in human, and 4-fold upregulation in dcKO mouse. *TGFB1I1* was 8-fold upregulated in human and 2.9-fold upregulated in mouse. *TGFB1I1* is thought to mediate progesterone resistance in endometriosis (46, 47). *GDF10* was 8.4-fold upregulated in human and 2.3-fold upregulated in mice (Figure 6C-D). Other TGFβ signaling genes that are predicted targets of other downregulated miRNAs were upregulated (Supplementary Figure S9).

**Figure 6.**
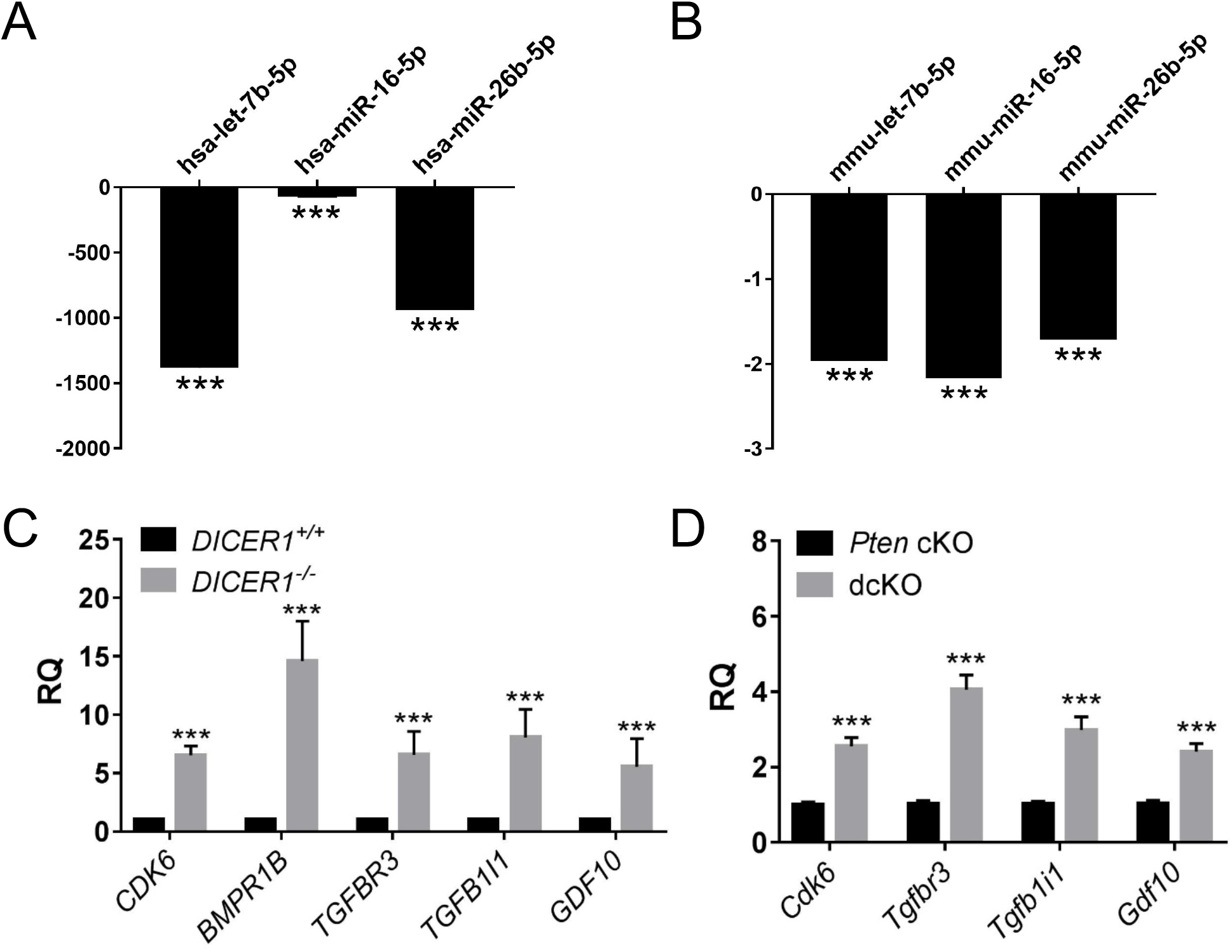
Downregulated miRNAs and upregulated TGFβ signaling genes. QPCR showed downregulation of miRNAs in human *DICER1^−/−^* cells (A) and dcKO mice (B). Mean ± SEM; two-tailed, Student’s *t*-test; ***, *P* < 0.001; *n* = 6; relative quantity of each miRNA by qPCR relative to endogenous control miRNA U6 snRNA (human) and snoRNA202 (mouse) normalized to *DICER1^+/+^* (A) or *Pten* cKO (B). QPCR showed upregulation of TGFβ signaling genes in human (C) and mouse (D). Mean ± SEM; ***, *P* < 0.001; Student’s t-test; *n*=6. RQ, relative quantity of each gene transcript by qPCR relative to endogenous control gene *GAPHD* (human) and 18S (mouse) normalized to *DICER1^+/+^* (C) or *Pten* cKO (D).

## Discussion

Endometrial cancer incidence continues to rise in the United States (1). Importantly, studies have shown that the increase in incidence is mostly in non-endometrioid or high-grade histologies, including serous and clear-cell adenocarcinomas (48). Although nearly all common cancers have improved cancer survival since the 1970s, endometrial cancer is one of the few cancers with an increased death rate (1). The highest risk factor linked to disease recurrence and death is a high tumor grade (5, 6). New therapies are needed for high-grade tumors, as these are the tumors leading to the most substantial mortality (3, 4).

In the present study, we describe the first mouse model of poorly-differentiated endometrial adenocarcinoma. Because *DICER1* mutations are frequently found in ultra-mutant tumors, *DICER1* mutations in endometrial adenocarcinoma were thought to be passenger mutations. However, recent genomic profiling of tumors through TCGA PanCancer and MSK-IMPACT showed enrichment of biallelic *DICER1* hotspot mutations in endometrial but not other cancers (8). Importantly, this analysis removed hypermutated tumors and the POLE subclass of endometrial cancers (8). *DICER1* mutants play a functional role in endometrial cancers, by affecting miRNA processing, miRNA expression, and miRNA-target gene expression (8, 11–15). These studies in large, robust, human endometrial cancer datasets show that *DICER1* mutations are not passenger mutations in endometrial cancer and suggest that DICER1 may play a unique yet significant role in endometrial cancer. Furthermore, on the molecular level, deregulated miRNA-target genes for let-7-5p and miR-16-5p family members were significantly enriched in the published *DICER1*-mutant TCGA datasets (8) and the present study (both dcKO mouse and human Ishikawa *DICER1^−/−^* cells; Figure 6). Thus, the dcKO mouse model and human Ishikawa *DICER1^−/−^* cells represent practical models.

Clinically, miR-16-5p and let-7-5p represent important molecular targets for cancer therapy. Treatment with liposomal miR-16 is being tested for mesothelioma (49) and is frequently considered a tumor suppressor (50). Let-7b-5p is a likely tumor suppressor in endometrial cancer (51). Also, targets of these miRNA molecules may be very important clinically. For example, CDK6 is a predicted target for multiple robustly downregulated miRNA molecules (Supplementary Table S5). Recent studies suggest that high expression of CDK6 may be a biomarker for poor prognosis endometrial cancers (52). Specifically, the high expression of CDK6 with high Ki67 expression was associated with shorter progression-free survival in women with endometrial cancer (52). Intact dcKO uterine tumors showed a significantly high frequency of Ki67-staining cells (Figure 5D). The use of CDK6 inhibitors, such as palbociclib, has shown promise *in vitro* and in orthotopic and xenograft models, particularly when tumors are *PTEN* mutant (53). Additionally, high expression of LIMK2 (Figure 5) suggests activation of ROCK/Rho signaling. Studies indicate that the use of T56-LIMKi, a LIM kinase inhibitor, leads to decreased proliferation and migration in cancer cells (39). Further, pre-clinical studies of a liposomal delivery system for EPHA2 knockdown have shown significant promise (54). Thus, the dcKO mouse and human Ishikawa *DICER1^−/−^* cells express target genes that recapitulate human cancers. Future studies will focus on testing these precision therapies in our mouse and cell models.

Poorly-differentiation endometrial tumors frequently develop within an atrophic, or postmenopausal endometrium (22). *Pgr^Cre/+;^ Dicer1^flox/flox^* mice have a benign single layer of luminal epithelium, lack of glandular epithelium, and thin to no endometrial stroma, a uterine phenotype consistent with atrophic endometrium from postmenopausal women (19). Additional deletion of *Pten* in this background of *Dicer1* deletion leads to poorly-differentiated adenocarcinoma. While hyper-proliferation of endometrial epithelium from endogenous or exogenous estradiol with mutation of *PTEN* is a biologically plausible mechanism for the development of endometrial cancer (22), little is known about the mechanism of development of endometrial cancer from atrophic endometrium. In most dcKO mice, the tumors seemed to arise from individual areas of the uterus rather than homogenously along the entire uterus. This mechanism is consistent with type II endometrial tumors primarily derived from heterogeneous endometrial polyps in women (22). The reason for the heterogeneous nature of the tumor development along the uterine horns is unknown. We suspect that a small portion of cells, potentially a hormone-independent stem cell population, maybe playing a role as decreased expression of *DICER1* in human endometrial cancers is associated with increased expression of stem cell markers (16). These studies will be the focus of future investigations.

Single allele deletions in *Dicer1* provide more aggressive disease in certain cancers when combined with oncogenes or deletion of tumors suppressors, while deletion of two alleles of *Dicer1* is protective (55–58). Conditional loss of one allele of *Dicer1* had a faster rate of lung cancer formation on an oncogenic *K-ras* background. In contrast, the loss of both *Dicer1* alleles led to inhibition of tumor formation (55). Similarly, *Dicer1* haploinsufficiency on an oncogenic *Braf(V600E)* background leads to increased metastasis in sarcomas (58). With the inactivation of the retinoblastoma gene, the loss of one allele of *Dicer1* in the retinoblasts of mice led to aggressive retinoblastoma (56). This effect seems to be oncogene dependent, as *Dicer1* haploinsufficiency on an oncogenic *c-Myc* background does not facilitate cancers (59). Thus, we initially hypothesized that aggressive endometrial carcinomas would arise from the loss of one allele of *Dicer1*, and the loss of two alleles of *Dicer1* would be protective. The survival curves and transcriptomic data suggest that deletion of a single allele of *Dicer* in a background of *Pten* deletion in the uterus does not lead to worse outcomes or even significant changes in gene expression compared to *Pten* deletion alone. Thus, we believe that the effects of a single allele loss of *Dicer1* are likely tissue-specific.

Further studies are needed on *Dicer1* in the adnexa. We believe that *Pten* cKO, *Pten-Dicer* het, and dcKO mice have metastatic endometrial cancer to the adnexa. There are a number of reasons for this conclusion: 1) the frequency of tumors in the adnexa is of lower penetrance than the uterus; 2) the histology of the adnexal tumors is similar to the uterine tumors; 3) the location of cancer in the adnexa (*i.e.*, ovarian hilum or adjacent to normal) suggests invasion or direct spread; 4) the large endometrial tumors with myometrial invasion; 5) multinodular location of adenocarcinoma along the oviduct with some areas of normal oviduct; and 6) no evidence of typical spread of oviductal or ovarian cancer such as peritoneal metastasis or ascites.

The poorly-differentiated adenocarcinomas of dcKO mice also exhibit features of clear-cell adenocarcinoma. Clinically, clear-cell endometrial adenocarcinomas are challenging to distinguish from high-grade endometrioid endometrial adenocarcinomas with clear-cell changes (28). In our mouse model, dcKO uteri show high-intensity staining for Napsin A, which is the most sensitive marker for clear-cell carcinoma (29). Additionally, dcKO tumors are HNF1B positive, another important marker of clear cells (28, 29). The use of the dcKO model as a model for clear-cell adenocarcinoma would be improved with comparison to transcriptomic studies from clear-cell endometrial adenocarcinoma from women. However, the only existing dataset of clear-cell endometrial cancer profiled five samples, used older microarray technology, and profiled a limited number of genes (60). We are encouraged that our gene sets were enriched in let-7 and miR-16 target genes similar to derepressed gene sets from *DICER1*-mutant TCGA tumors (8). Further studies are needed to transcriptomically interrogate pathologically well-characterized clear-cell endometrial adenocarcinoma from women for comparison to dcKO tumors.

Our data, as presented, show that the dcKO mouse represents poorly-differentiated adenocarcinoma. Mouse models of endometrial cancer that recapitulate human disease represent translational tools for a better understanding of the aggressive disease. We anticipate that this model will be highly relevant, not only to study the molecular characteristics of the rarer forms of human endometrial cancer but also to study the preclinical development of therapeutics.

## Materials and Methods

### Generation of mice and genotyping

All experiments conducted with mice were approved by the Indiana University School of Medicine Institutional Animal Care and Use Committee following the National Institutes of Health Guide for the Care and Use of Laboratory Animals. All animals were bred and kept under standard conditions. *Pgr^cre/+^*; *Dicer1^flox/flox^* (19) mice were bred to *Pgr^+/+^*; *Pten^flox/flox^* mice (21) and maintained on a C57BL/6J;129S5/Brd mixed hybrid background. Crossbreeding was used to generate experimental female mice. Mice were genotyped at 12-14 days of postnatal life and postmortem from tail biopsies (19, 21) (Supplementary Methods and Supplementary Figure S10).

### Survival studies, tissue collection, histological analysis, immunohistochemistry, and immunofluorescence staining

For survival studies, mice were caged and examined twice weekly. Mice were euthanized at humane endpoints of loss of normal behavior or ambulation, obvious distress, not eating or drinking, loss of 20% of body weight, tumor size greater than 10% of body weight, ulcerations, roughened hair coat or hunchback, or poor body conditioning score (61). At the time points listed, mice were sacrificed; body and uterine weight were recorded. One uterine horn, oviduct, and ovary were snap-frozen. The other uterine horn, oviduct, and ovary were fixed, and processing, paraffin embedding, and histological and immunohistochemistry staining were as described (19, 62). All histology was interpreted by clinical pathologists (R.E.E. and R.R.B.), with three sections each of 6-25 mice represented. FIGO tumor grading was used (Supplementary Table S6). For simplification of analyses, a binary FIGO grading of low-grade defined as FIGO grade 1 or 2 and high-grade defined as FIGO grade 3, was used as clinically recommended (24). Primary antibodies and conditions are listed (Supplementary Table S6). Immunohistochemistry staining was imaged on a Zeiss Axio Lab.A1 microscope (Oberkochen, Germany) and scored based on the frequency of staining as the percentage of positive staining cells under low-power (5X) fields and confirmed with ImageJ using FIJI (63–66). Immunofluorescence staining was imaged on an EVOS FL Cell Imaging System (Thermo Fisher Scientific, Waltham, MA).

### Removal of ovaries and steroid hormone treatment

Mice underwent ovariectomy (ovex) at six weeks. Mice were randomly divided into treatment groups: progesterone pellet (Innovative Research of America, Sarasota, Florida, 25mg/60-day release pellet) or placebo pellet, and uterine tissues collected after 60 days.

### *In vitro* growth conditions, CRISPR-Cas9 *DICER1* deletion, and characterization

Ishikawa cells were obtained from the Cytogenetics and Cell Authentication Core at MD Anderson Cancer Center (Houston, TX) and maintained in RPMI1640 (Thermo Fisher Scientific) supplemented with 10% fetal bovine serum (Atlanta Biologicals, Minneapolis, MN) and 1% penicillin/streptomycin (Thermo Fisher Scientific). Cell line authentication was confirmed using a short tandem repeat (STR) marker profile and interspecies contamination check (IDEXX BioAnalytics, Westbrook, ME). *DICER1* was deleted in Ishikawa cells using CRISPR-Cas9 (67) (Supplementary Methods and Supplementary Figure S11). Briefly, Ishikawa cells were transfected using lipofectamine 2000 (Thermo Fisher Scientific) with pX330-U6-Chimeric_BB-CBh-hSpCas9 containing a guide RNA (gRNA) to human *DICER1* [Cell-based Assay Screening Service (C-BASS) core, Baylor College of Medicine (Houston, TX)]. Single-cell colonies were grown and screened by end-point PCR. Positive clones underwent Sanger sequencing (Genewiz, South Plainfield, NJ) to identify the location of DNA breaks. Preparation of total cell lysates, western blot, proliferation, and colony formation assay are described in Supplementary Methods. Supplementary Table S6 lists the antibodies and conditions.

### RNA isolation, next-generation miRNA sequencing, whole-genome expression analysis, and real-time quantitative PCR analysis

Total RNA was extracted from uteri and Ishikawa cells, DNase treated, and assessed for quality (19). Details of next-generation sequencing are provided in Supplementary Methods. Briefly, poly-A RNA libraries were constructed using mRNA Stranded TruSeq protocol (Illumina, San Diego, CA). Small RNA library construction was performed using the TruSeq Small RNA kit (Illumina). Purified libraries were visualized and quantified using a TapeStation HSD1000 (Agilent, Santa Clara, CA). Sequencing was performed on an Illumina NextSeq500 instrument at the Center for Genomics and Bioinformatics (Bloomington, IN). The mRNA reads were mapped against GRCm38 and GRCh38.p5. Initial mRNA differential expression analysis was carried out using the DESeq2 package (version 1.10.1) in R/Bioconductor (R version 3.2.1) (68). MiRNA reads were mapped to miRBase V21. Initial miRNA differential expression analysis was carried out as described (69). Dysregulated canonical pathways were determined from Ingenuity Pathway Analysis (Qiagen Bioinformatics, Redwood City, CA). Integrated miRNA:mRNA analyses were carried out using miRTarBase (70). Reverse transcription and real-time qPCR were performed on a QuantStudio3 Real-Time PCR System (Thermo Fisher Scientific) and analyzed (19). Supplementary Table S6 lists the TaqMan assays and Sybr primer sequences.

### Statistical analysis

Statistical analysis was performed with one of the following: Student’s *t*-test, Fisher’s exact test, Chi-squared test, multiple *t*-test, 2-way ANOVA, or log-rank (Mantel-Cox) pairwise comparison with Bonferroni correction. Differences between groups were determined by *P*<0.05. Statistical analyses were conducted using the InStat package for Prism8 (GraphPad, San Diego, CA).

## Acknowledgments

We acknowledge the Indiana Center for Musculoskeletal Health Histology Core at Indiana University School of Medicine and the Human Tissue and Acquisition and Pathology Core at the Dan L. Duncan Comprehensive Cancer Center at Baylor College of Medicine for histology services; the Center for Genomics and Bioinformatics for RNA sequencing, and the Center for Medical Genomics for RNA quality control analysis. We appreciate Dr. Ken Nephew for thoughtful review and comments.

