## Supplementary for "*Pten* and *Dicer1* loss causes poorly-differentiated endometrial adenocarcinoma in mice"

[a]Department of Obstetrics and Gynecology, Indiana University School of Medicine, Indianapolis, IN 46202; [b]Department of Pathology and Laboratory Medicine, Indiana University School of Medicine, Indianapolis, IN 46202; [c]Department of Pathology and Laboratory Medicine, University of North Carolina School of Medicine, Chapel Hill, NC 27599; [d]Department of Medicine, Baylor College of Medicine, Houston, TX 77030; [e]Center for Genomics and Bioinformatics, Indiana University, Bloomington, IN 47405; [f]National Institute of Environmental Health Sciences, Research Triangle Park, NC 27709; [g]Department of Molecular and Cellular Biology, Baylor College of Medicine, Houston, TX 77030

[1]To whom correspondence should be addressed
Shannon M. Hawkins, M.D., Ph.D.
Indiana University School of Medicine at
Indiana University Purdue University at Indianapolis (IUPUI)
550 N. University Blvd, UH2440
Indianapolis, IN 46202
0000-0002-0727-3971

**This PDF file includes:**

Supplementary text
Figures S1 to S11
SI References

**Other supplementary materials for this manuscript include the following:**

Datasets S1 to S6

**Supplementary Methods:**
**Genotyping.**

Following genomic DNA isolation from tail snips (1), PCR was used to detect the *floxed Pten* and/or *Dicer* and *Cre* alleles using primers in Supplementary Figure S1. For the *Cre* allele, the following conditions were used with *Pgr Cre* P1, P2, and P3 primers: 10 minutes at 95°C, followed by 35 cycles of 30 seconds at 95°C (denaturation), 30 seconds at 63°C (annealing), and 30 seconds at 72°C (extension) with final extension at 72°C for 10 minutes. For the *Pten* allele, the following conditions were used with *Pten* F and R primers: 10 minutes at 95°C, followed by 35 cycles of 30 seconds at 95°C (denaturation), 60 seconds at 60°C (annealing), and 60 seconds at 72°C (extension) with final extension at 72°C for 10 minutes. For the *Dicer* allele, the following conditions were used with *Dicer* D1, D2, and D3 primers: 10 minutes at 95°C, followed by 35 cycles of 30 seconds at 95°C (denaturation), 45 seconds at 60°C (annealing), and 45 seconds at 72°C (extension) with final extension at 72°C for 10 minutes (2, 3). The PCR amplicons were separated using a 2% agarose gel, and the results were visualized using ChemiDoc™ MP Imaging System (Bio-Rad; Hercules, CA).

***DICER1* deletion using CRISPR-Cas9.**

Using CRISPR-Cas9 gene editing technology (4), we generated cell lines – *DICER1*$^{+/-}$ and *DICER1*$^{-/-}$. One day prior to transfection, Ishikawa cells were seeded in a 6-well plate at 200,000 cells per well. Lipofectamine 2000 (Thermo Fisher Scientific; Waltham, MA) transfection was carried out per manufacturer's instructions using 1-µg of each guide RNA (gRNA) in pX330-U6-Chimeric_BB-

CBh-hSpCas9. Supplementary Figure S11 depicts the relative location of gRNA1 and gRNA2 to human *DICER1* and gives the sequence of each gRNA. After incubation for 48 hours in a humidified incubator at 37°C and 5% of $CO_2$, 1 cell per well was plated into a 96-well plate. Colonies were expanded into 6-well plates. Genomic DNA was extracted with lysis buffer [10 mM Tris-HCl, pH 8.5; 5 mM EDTA; 200 mM NaCl; 0.2% SDS with 1 mg/ml Proteinase K (DOT Scientific; Burton, MI)], followed by sequential 5M NaCl protein precipitation and isopropanol DNA precipitation. Genomic DNA was solubilized in nuclease-free water (Thermo Fisher Scientific). Deletion screening was performed by end-point PCR amplification. The following conditions were used with DICER1-F and -R primers: 5 minutes at 95°C, followed by 35 cycles of 30 seconds at 95°C (denaturation), 45 seconds at 62°C (annealing), and 30 seconds at 72°C (extension) with final extension at 72°C for 10 minutes. The amplicons were visualized using a ChemiDoc™ MP Imaging System (Bio-Rad). DICER1-F and -R primers were used for Sanger sequencing (Genewiz; South Plainfield, NJ) in both directions (Entrez Gene ID23405). FinchTV (Geospiza, Inc., Seatle, WA) was used to view DNA sequencing.

**Western blot.**

Protein extracts were prepared from the positive clones in RIPA (radioimmunoprecipitation assay) buffer (Thermo Fisher Scientific). BCA (bicinchoninic acid) Protein Assay Reagent Kit (Thermo Fisher Scientific) was used to quantify lysed proteins. 30-μg of protein lysates were electrophoresed through a NuPAGE Tris-Acetate Mini Gel (Thermo Fisher Scientific) and then transferred to a polyvinylidene difluoride (PVDF) membrane (Thermo Fisher Scientific). Immunoblotting was performed using a rabbit polyclonal antibody against DICER1 at a 1:200 dilution (Sigma-Aldrich; St. Louis, MO) and a mouse monoclonal antibody against β-actin at a 1:1000 dilution (Santa Cruz Biotechnology; Dallas, TX). Bands were visualized using the SuperSignal West Pico Chemiluminescent Substrate (Thermo Fisher Scientific) on a ChemiDoc™ MP Imaging System (Bio-Rad).

**Proliferation.**

Ishikawa cells were seeded in 96-well plates at a density of 1,000 cells/well in triplicate. CellTiter 96 Aqueous One Solution Cell Proliferation assay (Promega, Madison, WI) was used. The optical density at 496 nm and 630 nm were measured using a Synergy H1 Hybrid Multi-Mode Microplate Reader (BioTek; Winooski, VT) for 1, 2, 3, and 4 days. After subtraction of background (media only), cell proliferation was presented relative to each cell line on day 1.

**Colony formation assay.**

A total of 500 Ishikawa cells were seeded in a 6-well plate in triplicate for 10-14 days or until they formed sufficient clones consisting of 50 or more cells. The cells were fixed with methanol (Thermo Fisher Scientific) for 15 minutes at room temperature. Cells were stained with 0.1% crystal violet (Sigma-Aldrich) for 10 minutes at room temperature. After rinsing, colonies that had formed in each well were counted under microscopy (Nikon Eclipse TS100-F Inverted Microscope; Minato City, Tokyo, Japan).

**RNA analysis**

NextSeq read sequences for both mouse and human were cleaned using Trimmomatic version 0.32 with the parameters “2:20:5 LEADING:3 TRAILING:3 SLIDINGWINDOW:4:15 MINLEN:18” to remove adapter sequences and perform quality trimming (5). Custom Perl scripts along with FASTX-Toolkit version 0.0.13.2 with the following parameters, “-l 17 -v -M 1 –I,” were also used to look for partial adapters within the reads (6). The mRNA reads were mapped against GRCm38 or GRCh38.p5 using the “—strata” parameter and reporting up to 50 alignments per read (7) and using TopHat2 version 2.1.1 with parameters “--b2-very-sensitive --read-edit-dist 2 --max-multihits 100” (8). The multicov option within Bedtools was used to produce counts for each gene, allowing for multi-mapped reads to be counted for each feature they are associated with (9). Read counts for each gene were created using HTSeq-Count from the HTSeq package version 0.6.1p1and Gencode v24 as the annotation (10, 11). Custom Perl scripts were used for estimation of transcript abundances based on Fragments Per Kilobase of exon per Million fragments mapped (FPKM). To detect known mature microRNA sequences, MiRDeep2 version 2.0.0.7 was run using the current version of miRBase version 21 as reference (12, 13). MiRDeep2 uses bowtie, version 1.1.2, to perform mapping of the reads and includes tools for the identification and quantification of microRNAs (7). Multiple testing correction at a false discovery rate $< 0.05$ was applied to identify differentially expressed genes. The “ggplot2” library package in R was used to make display figures, including volcano and principal component analysis plots.

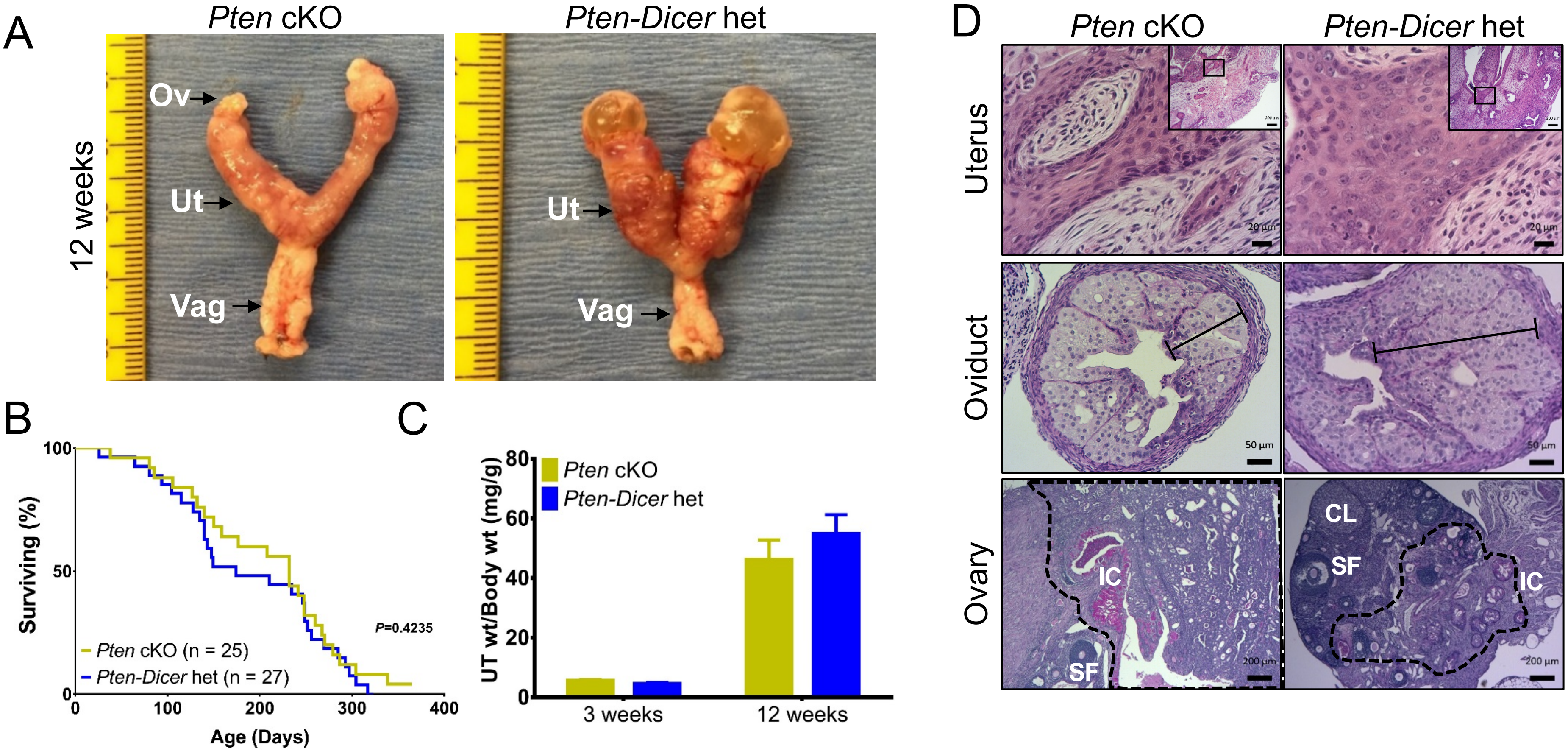


**Supplementary Figure S1.** Similar malignant phenotype between *Pten* cKO and *Pten-Dicer* het mice. (A) *Pten-Dicer* het uteri showed similar gross morphology as *Pten* cKO uteri. By 12 weeks, *Pten* cKO and *Pten-Dicer* het uteri had enlarged uterine horns. Ov, ovary; Ut, uterus; Vag, vagina. Ruler, tick marks indicate 1 mm. (B) Kaplan-Meier survival curve, analyzed by log-rank (Mantel-Cox) pairwise comparison with Bonferroni correction, showed no survival difference between *Pten* cKO and *Pten-Dicer* het mice. The median survival for *Pten-Dicer* het mice was 198 days. (C) There was no statistical difference in uterine weight between *Pten* cKO and *Pten-Dicer* het at 3 and 12 weeks of age. Mean ± SEM; Student's *t*-test, $n > 8$. (D) Similar endometrial adenocarcinoma (top), atypical epithelial cells in oviduct (middle), and invasive adenocarcinoma (IC) into the hilum of ovaries (bottom) of both *Pten* cKO and *Pten-Dicer* het mice. Hematoxylin and eosin (H&E) staining of uteri and Periodic acid-Schiff (PAS) staining of oviduct and ovary. Line, multiple cell layers of atypical oviductal epithelium; CL, corpus luteum; SF, secondary follicle; IC, invasive cancer; Black dashed lines, well-differentiated adenocarcinoma invading at ovarian hilum. (Scale bars, low-power uteri, 200 µm; high-power uteri, 20 µm; oviduct, 50 µm; ovary, 200 µm.)

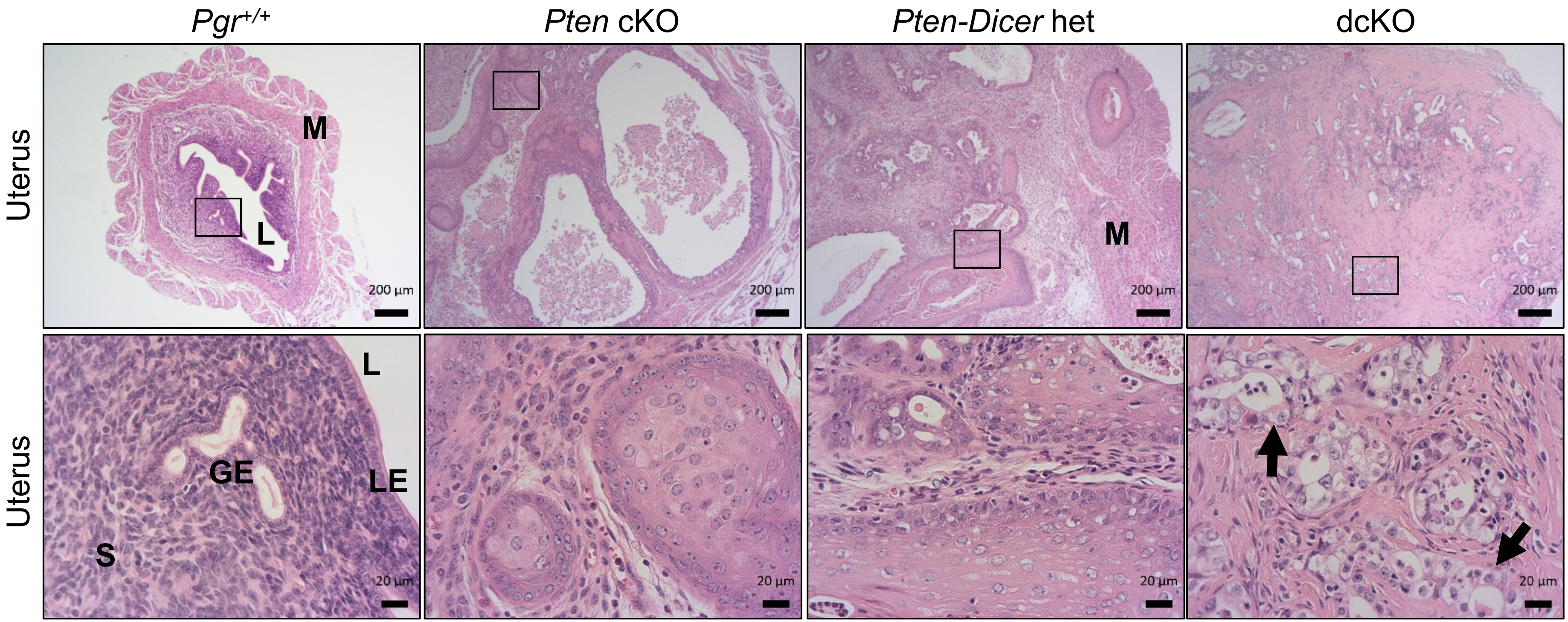


**Supplementary Figure S2.** Histology of 6-month-old uteri. *Pgr*$^{+/+}$ uteri showed normal histology that included two distinct layers of myometrium (M), a single layer of luminal epithelium (LE) surrounding the lumen (L), and normal appearing glandular epithelium (GE) and endometrial stroma (S). *Pten* cKO and *Pten-Dicer* het uteri showed well-differentiated adenocarcinoma, with confluent growth of endometrial glands with squamous metaplasia, minimal atypia, and low mitotic activity. Uteri from dcKO mice showed poorly-differentiated adenocarcinoma with more solid structures lacking glandular features. Arrows indicate malignant epithelial cells with large, hyperchromatic nuclei and pale-staining to clear cytoplasm. H&E. (Scale bars, low-power, 200 µm; high-power, 20 µm.)

Supplementary Figure S3

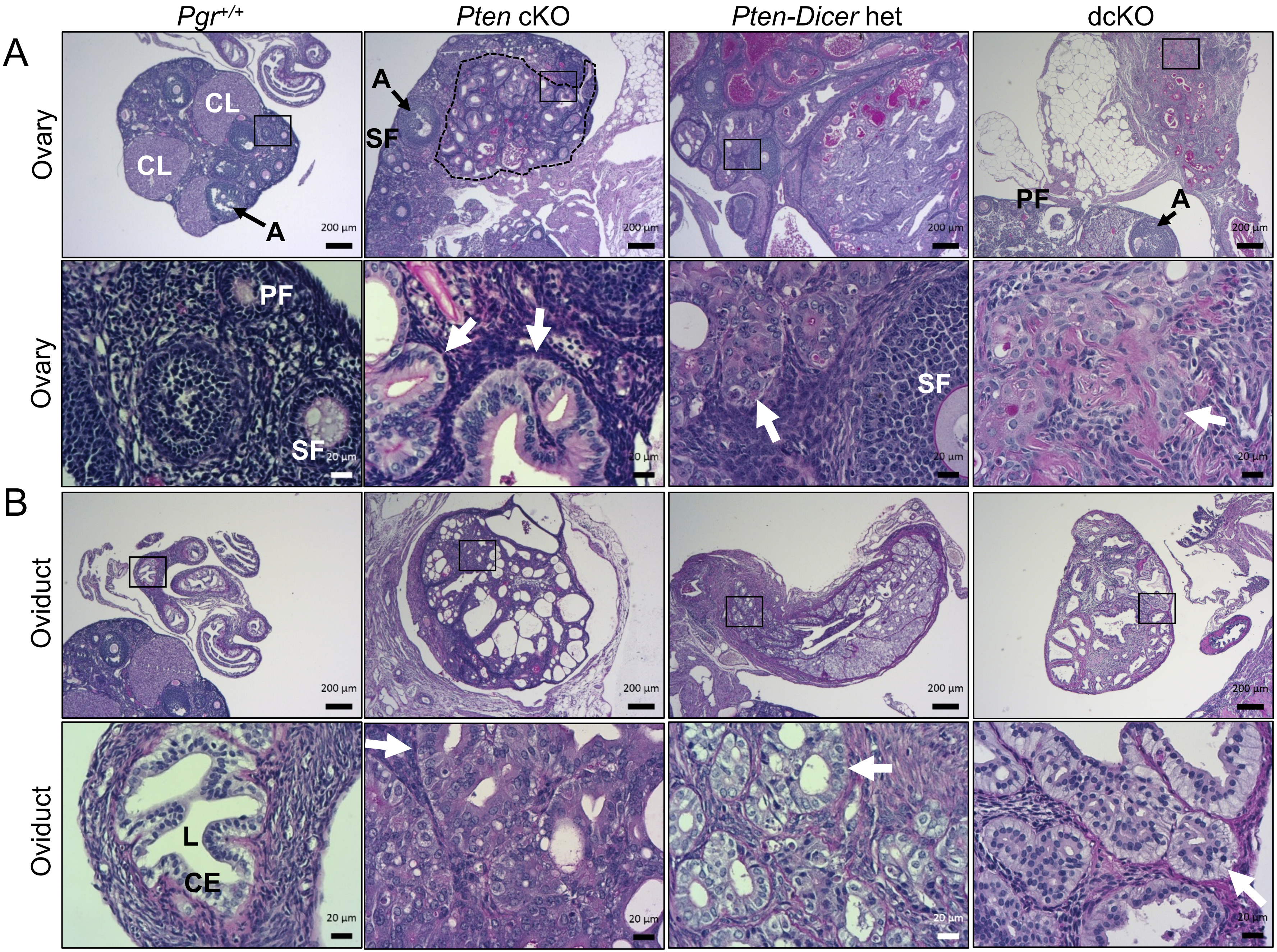

**Supplementary Figure S3.** Histology of 6-month-old adnexa. (A) *Pgr*$^{+/+}$ ovary showed normal histology with follicles in each stage of follicular development. There were primary (PF), secondary (SF), and antral (A) follicles and corpus luteum (CL). *Pten* cKO ovaries contained normal appearing follicles including secondary and antral follicles. However, there was also well-differentiated adenocarcinoma (dotted lines) that appeared to be invading at the ovarian hilum. The *Pten-Dicer* het ovary section had a secondary follicle, but the ovary is encompassed by well-differentiated adenocarcinoma. Ovaries from dcKO mice showed normal follicles as evidenced by primary and antral follicles, and the adenocarcinoma was clearly distinct from the ovary, being found in the fat tissue surrounding the ovary (box, higher magnification, white arrow). (B) *Pgr*$^{+/+}$ oviduct showed normal histology with a single layer of columnar epithelium (CE) surrounded by a thin stroma and myometrial layer. A lumen (L) was noted. *Pten* cKO, *Pten-Dicer* het, and dcKO oviducts were filled with adenocarcinoma. PAS. White arrows indicate adenocarcinoma. (Scale bars, low-power, 200 μm; high-power, 20 μm.)

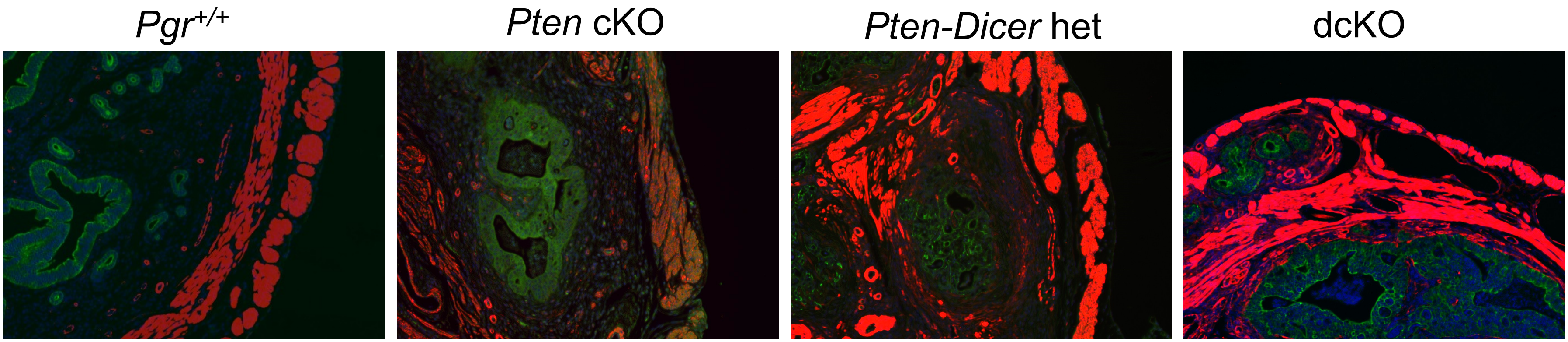


**Supplementary Figure S4.** Invasive uterine adenocarcinoma at 12 weeks. Immunofluorescence staining showed cytokeratin-8 (green)-stained epithelium surrounded by smooth muscle actin (red)-stained myometrium in *Pgr*$^{+/+}$ uteri. *Pten* cKO, *Pten-Dicer* het, and dcKO uteri showed growth of epithelial cells between the layers of the myometrium. DAPI, blue. 4x magnification.

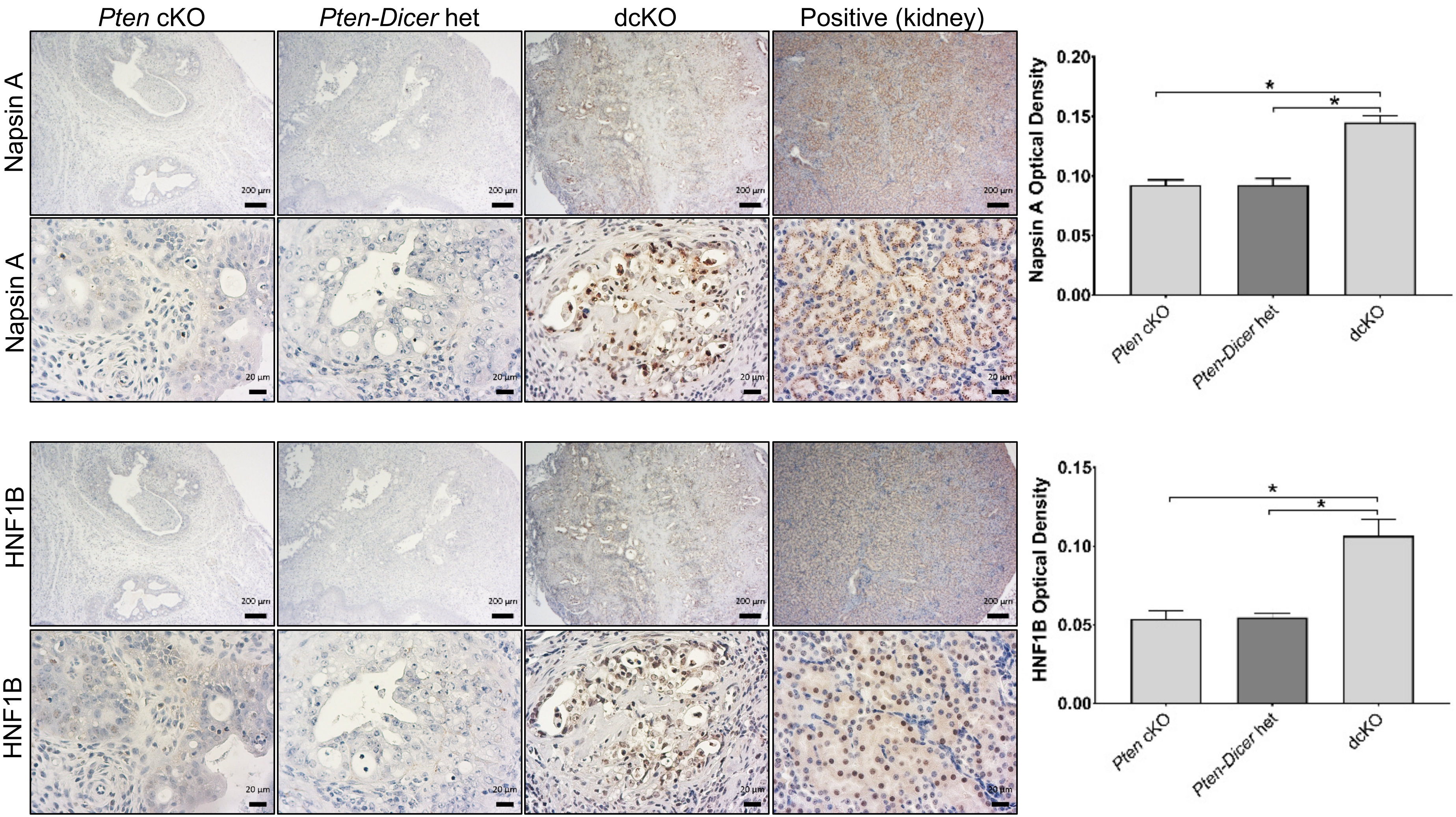


**Supplementary Figure S5.** Uteri from dcKO mice express clear-cell adenocarcinoma markers. Lower magnification views of Napsin A and HNF1B immunohistochemistry from Figure 3. Napsin A was cytoplasmic, and HNF1B was nuclear. Kidney is positive control. Confirmatory quantification using ImageJ with FIJI. Mean ± SEM; *, $P < 0.05$, multiple *t*-test, $n = 6$. (Scale bars, low-power, 200µm; high-power, 20 µm.)

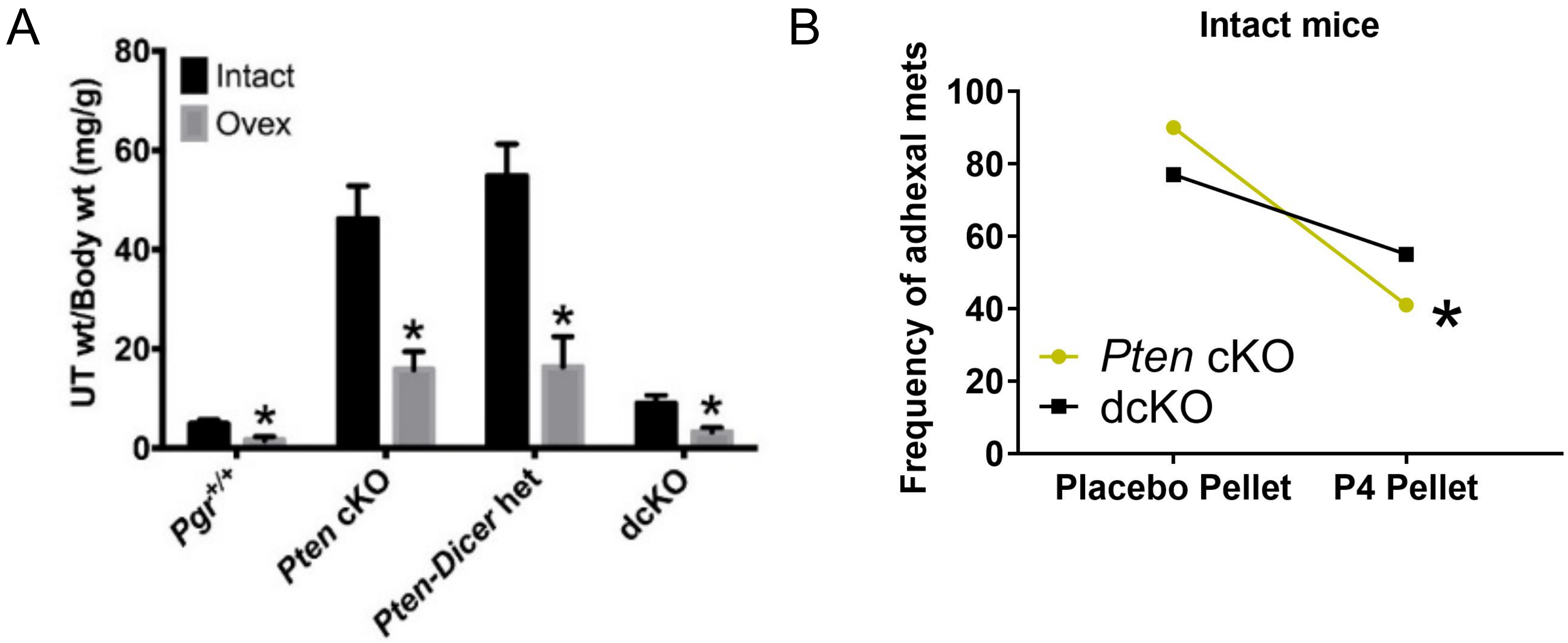


**Supplementary Figure S6.** The effects of ovarian insufficiency and long-term progesterone therapy. (A) After removal of ovaries (ovex) or not (intact) at 6 weeks, mice were dissected at 12 weeks, and uteri were weighed and normalized to body weight. Removal of ovaries resulted in smaller uteri for each genotype, but ovex *Pten* cKO and *Pten-Dicer* het uteri remained large. The frequency of poorly-differentiated endometrial adenocarcinomas was not significantly decreased with progesterone treatment (Supplementary Table S2). Mean ±SEM; *, $P < 0.05$, multiple *t*-test, $n > 6$. (B). Treatment with 60 days of progesterone significantly decreased the frequency of metastasis to the adnexa of *Pten* cKO mice. Fisher's exact test. *, $P<0.05$, $n > 9$.

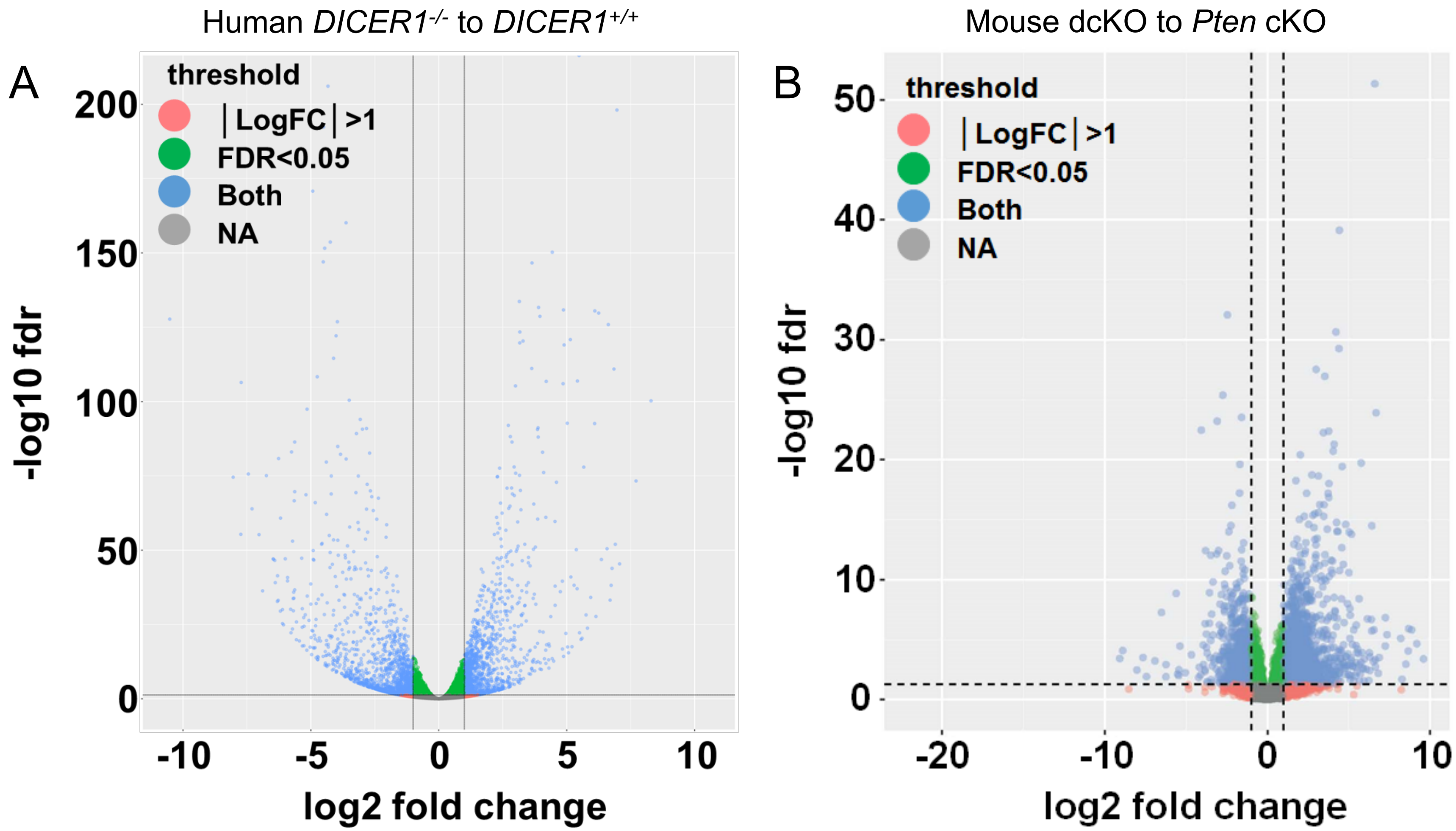


**Supplementary Figure S7:** Dysregulated genes with loss of two alleles of *DICER1*. Volcano plot using FDR<0.05 and log2 fold change >1 or <-1 for Ishikawa *DICER1*$^{-/-}$ cells (A) or dcKO mouse uteri (B).

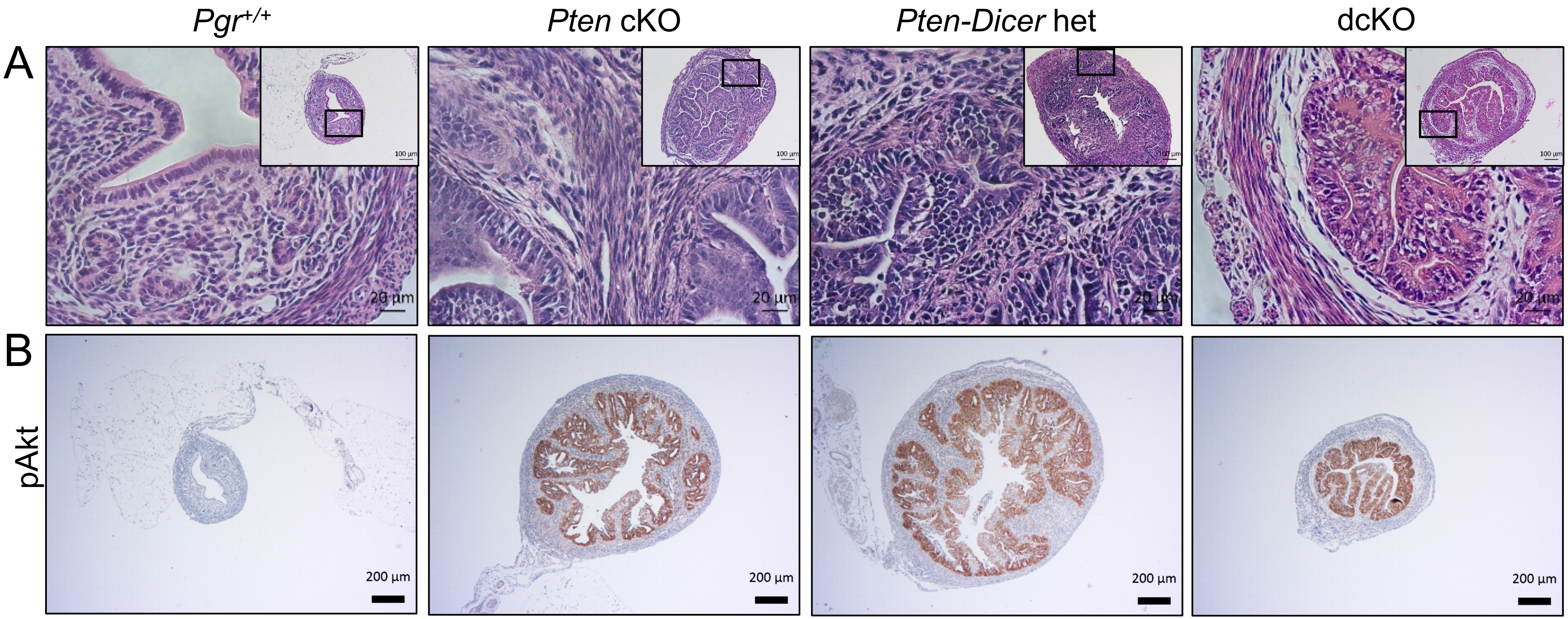


**Supplementary Figure S8:** Early adenocarcinoma with epithelial expression of phospho-AKT. (A) At three weeks, *Pten* cKO, *Pten-Dicer* het, and dcKO exhibited adenocarcinoma consisting of confluent growth of epithelium with pushing borders into the endometrial stromal and myometrium. (Scale bars, low-power, 100 µm; high-power, 20 µm.) (B) Significant phospho-AKT staining in endometrial epithelial cells in *Pten* cKO, *Pten-Dicer* het, and dcKO uteri at 3 weeks. (Scale bars, 200 µm.)

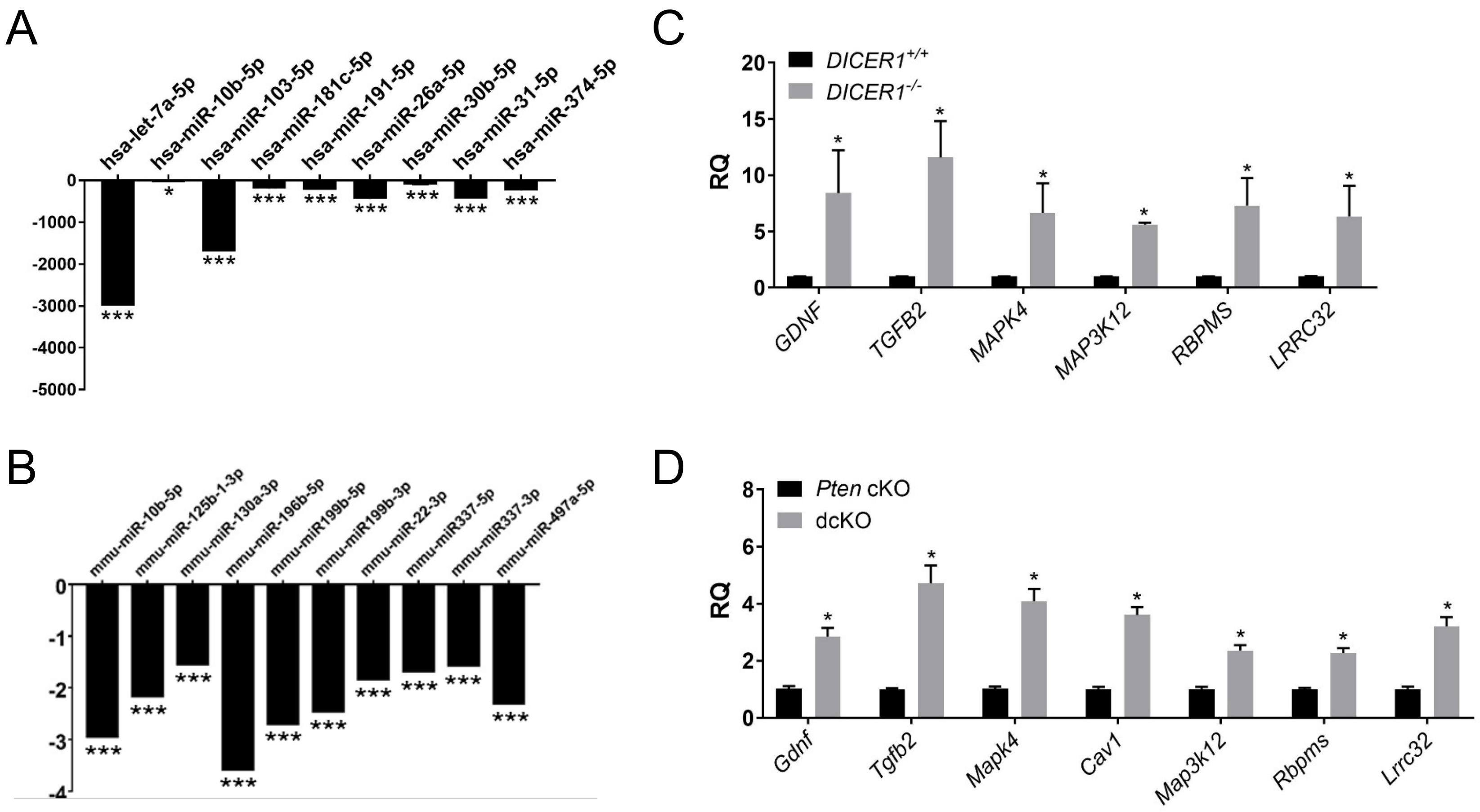


**Supplementary Figure S9.** Downregulation of miRNAs and upregulation of miRNA-target genes with *DICER1* deletion. QPCR showed downregulation of miRNAs in human *DICER1*$^{-/-}$ cells (A) and dcKO mice (B). Mean ± SEM; two-tailed Student's *t*-test; ***, $P < 0.001$; $n = 6$; relative quantity of each miRNA by qPCR relative to endogenous control miRNA U6 snRNA (human) and snoRNA202 (mouse) normalized to *DICER1*$^{+/+}$ (A) or *Pten* cKO (B). QPCR showed upregulation of TGFβ signaling genes in human *DICER1*$^{-/-}$ cells (C) and dcKO mice (D). Mean± SEM; two-tailed Student's *t*-test; *, $P < 0.05$; $n = 6$; RQ, relative quantity of each gene transcript relative to endogenous control gene *GAPHD* (human) and 18S (mouse) normalized to *DICER1*$^{+/+}$ (C) or *Pten* cKO (D).

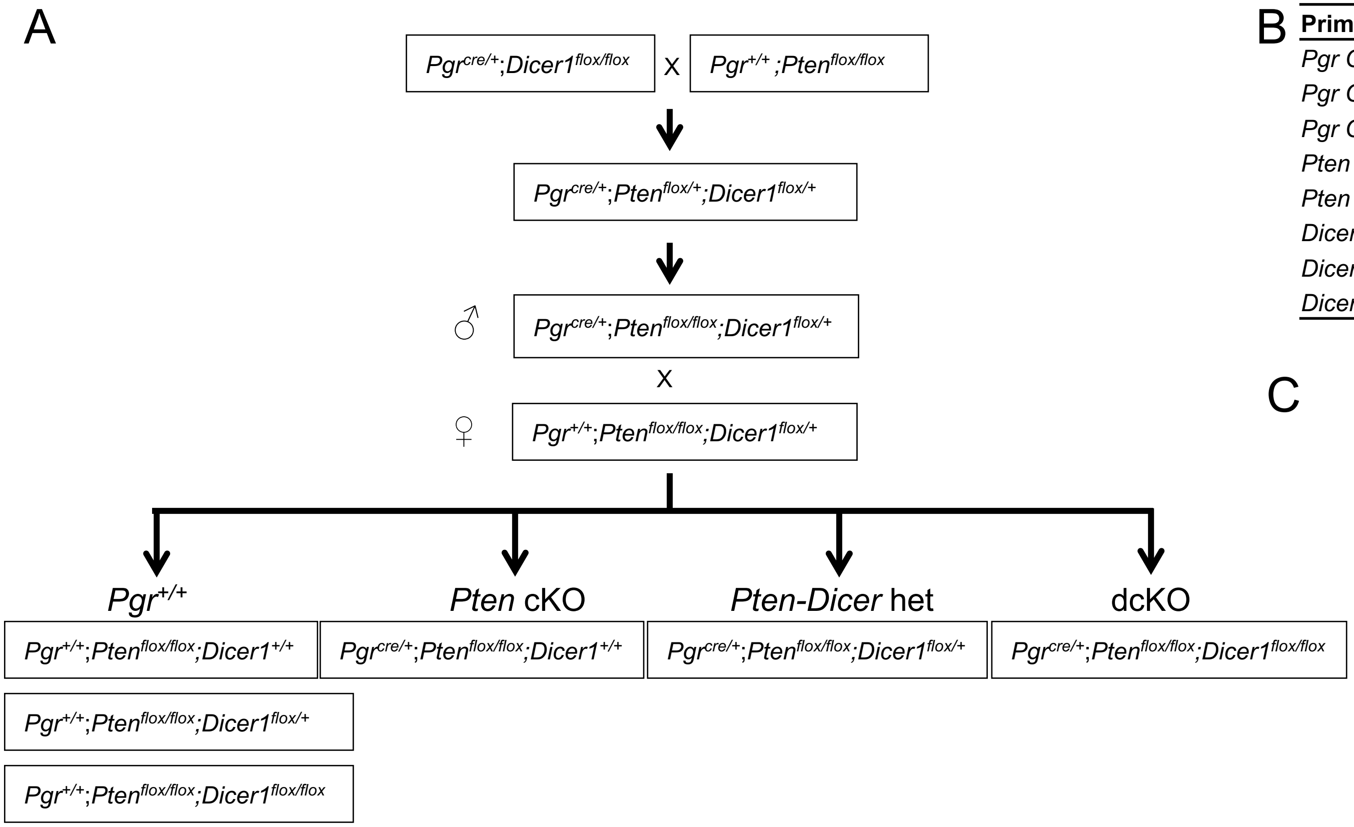


B

| Primer Name | Primer Sequence |
|---|---|
| *Pgr Cre* P1 | 5'- TATACCGATCTCCCTGGACG -3' |
| *Pgr Cre* P2 | 5'-ATGTTTAGCTGGCCCAAATG -3' |
| *Pgr Cre* P3 | 5'- CCCAAAGAGACACCAGGAAG -3' |
| *Pten* F | 5'-CAAGCACTCTGCGAACTGAG -3' |
| *Pten* R | 5'-AAGTTTTTGAAGGCAAGATGC -3' |
| *Dicer1* D1 | 5'-GGTTACATGGCTAGACTCAAAGC -3' |
| *Dicer1* D2 | 5'-AGGTGCCTTTCGTTTAGGAAC -3' |
| *Dicer1* D3 | 5'-AAAGCAGAACTCTAATGCCCC -3' |

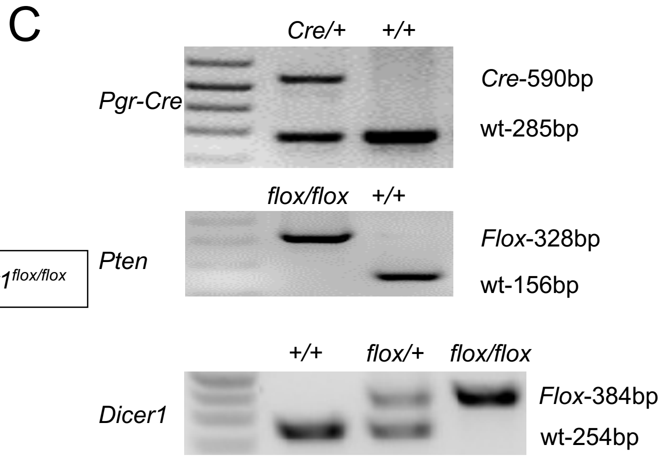


**Supplementary Figure S10.** Generation of mice with conditional deletion of *Pten* and *Dicer1* in the mouse uterus with *Pgr*$^{Cre}$. (A) Breeding strategy with *Pgr*$^{Cre}$ maintained on the male mice. (B) Primers for genotyping. Details of genotyping are described in Supplementary Methods. (C) Tail genotyping showing the *Cre*, *floxed*, or wild-type allele.

A

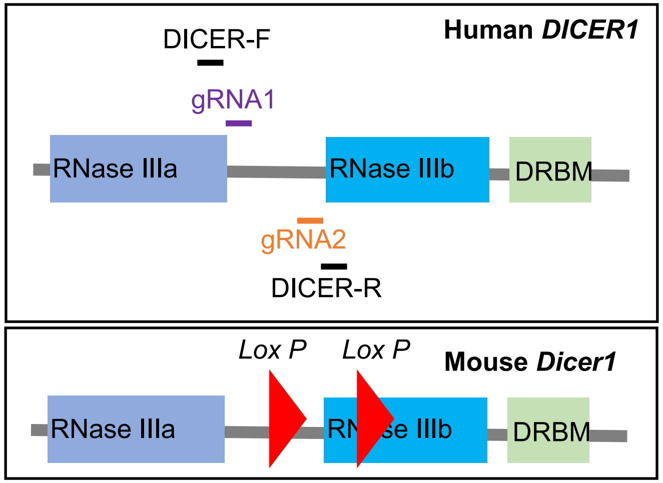


B

| | Name | Sequence | locations |
|---|---|---|---|
| sgRNAs | gRNA1 | 5’-TGACTATGAAGATGATTTCCTG-3’ | 4623-4644 bp |
| | gRNA2 | 5’-CACTGAATCACCTTATATCGGGG-3’ | 5288-5310 bp |
| PCR primers | DICER1-F | 5’-TGGCAAACTGGATGAGGATT-3’ | 4545-4565 bp |
| | DICER1-R | 5’-GTGGGCTCCTTACCAGTGAT-3’ | 5404-5424 bp |

C

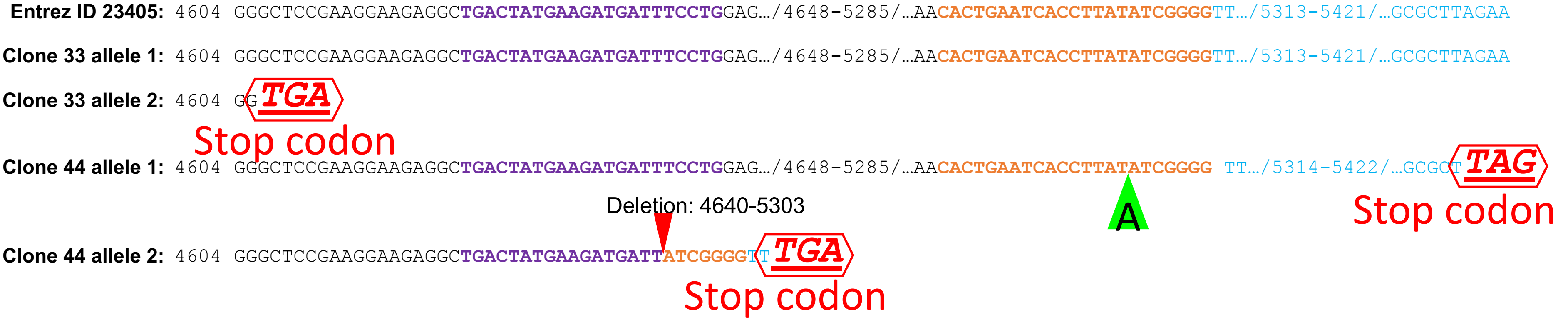


**Supplementary Figure S11.** Deletion of *DICER1* in Ishikawa cells using CRISPR-Cas9. (A) Depiction of human *DICER1* showing relative location of gRNA1 and gRNA2 to RNase IIIa, RNase IIIb, and the DRBM (double-stranded RNA binding motif) domains. Location of gRNA molecules were chosen based on best sequence that aligned with floxed alleles in mouse *Dicer1*. (B) Sequence of gRNA1 and gRNA2 that were cloned into pX330-U6-Chimeric_BB-CBh-hSpCas9. End-point PCR primers used for single-cell clone screening. (C) Genomic sequences of CRISPR-Cas9-edited Ishikawa clones revealed deletions leading to premature stop codons. Purple font is gRNA1; brown font is gRNA2; read arrow indicates deletion; green arrow indicates insertion; red stop sign indicates novel stop codon.

**Dataset S1 (separate file). Supplementary Table S1.** Female reproductive tract pathology for *Pten-Dicer* het mice.

**Dataset S2 (separate file**). **Supplementary Table S2.** Female reproductive tract pathology for *Pten* cKO and dcKO mice.

**Dataset S3 (separate file). Supplementary Table S3.** Differentially expressed genes and pathways in dcKO mice and cells.

**Dataset S4 (separate file). Supplementary Table S4.** Steroid hormone gene set enrichment analyses.

**Dataset S5 (separate file). Supplementary Table S5.** Differentially expressed miRNAs and miRNA-target genes and pathways in dcKO mice and cells.

**Dataset S6 (separate file). Supplementary Table S6.** Antibodies and primers.
